# Neurotensin receptor 2 is induced in astrocytes and brain endothelial cells in relation to status epilepticus and neuroinflammation following pilocarpine administration in rats

**DOI:** 10.1101/2020.06.29.166637

**Authors:** Kyriatzis Grigorios, Bernard Anne, Bôle Angélique, Pflieger Guillaume, Chalas Petros, Masse Maxime, Lécorché Pascaline, Jacquot Guillaume, Ferhat Lotfi, Khrestchatisky Michel

**Affiliations:** Aix-Marseille Univ, CNRS, INP, Inst Neurophysiopathol, Marseille, France; VECT-HORUS, Faculté de Médecine, 27 Bd Jean Moulin, 13385 Marseille Cedex 5, France

**Author notes:** Equally last and corresponding authors and. **MAIN POINTS** Epilepsy is associated with increased NTSR2 expression in astrocytes and endothelial cells. Proinflammatory factors induce NTSR2 in astroglia with immediate early gene response. A neurotensin analog down regulates NTSR2 and glial cell inflammation.

**Keywords:** NT, NTS, NTSR2, seizures, epilepsy, microglia, endothelial cells, blood-brain barrier, immediate early gene

## Abstract

Neurotensin (NT) acts as a primary neurotransmitter and neuromodulator in the CNS and has been involved in a number of CNS pathologies including epilepsy. NT mediates its central and peripheral effects by interacting with the NTSR1, NTSR2 and NTSR3 receptor subtypes. To date, little is known about the precise expression of the NT receptors in brain neural cells and their regulation in pathology. In the present work, we studied expression of the NTSR2 protein in the rat hippocampus using a model of temporal lobe epilepsy induced by pilocarpine and questioned whether NTSR2 was modulated in conditions of neuro-inflammation. This model is characterized by a rapid and intense inflammatory reaction with a pattern of reactive gliosis in the hippocampus. We show that NTSR2 protein is expressed in hippocampal astrocytes and its expression increases together with astrocyte reactivity following induction of status epilepticus. NTSR2 immunoreactivity is also increased in perivascular astrocytes and their end-feet and is apparent in endothelial cells following induction of status epilepticus. Proinflammatory factors such as IL1β and LPS induced NTSR2 in astrocytes, but also in microglia *in vitro*. Glial NTSR2 expression showed characteristic immediate early gene response under inflammatory conditions. Treating inflamed glial cells with a vectorized NT analogue decreased NTSR2 expression as well as astrocytic and microglial reactivity. Together, these results suggest that NTSR2 is implicated in astroglial and gliovascular inflammation and that targeting the NTSR2 receptor may open new avenues in the regulation of neuroinflammation in CNS diseases.

**TABLE OF CONTENTS IMAGE:** 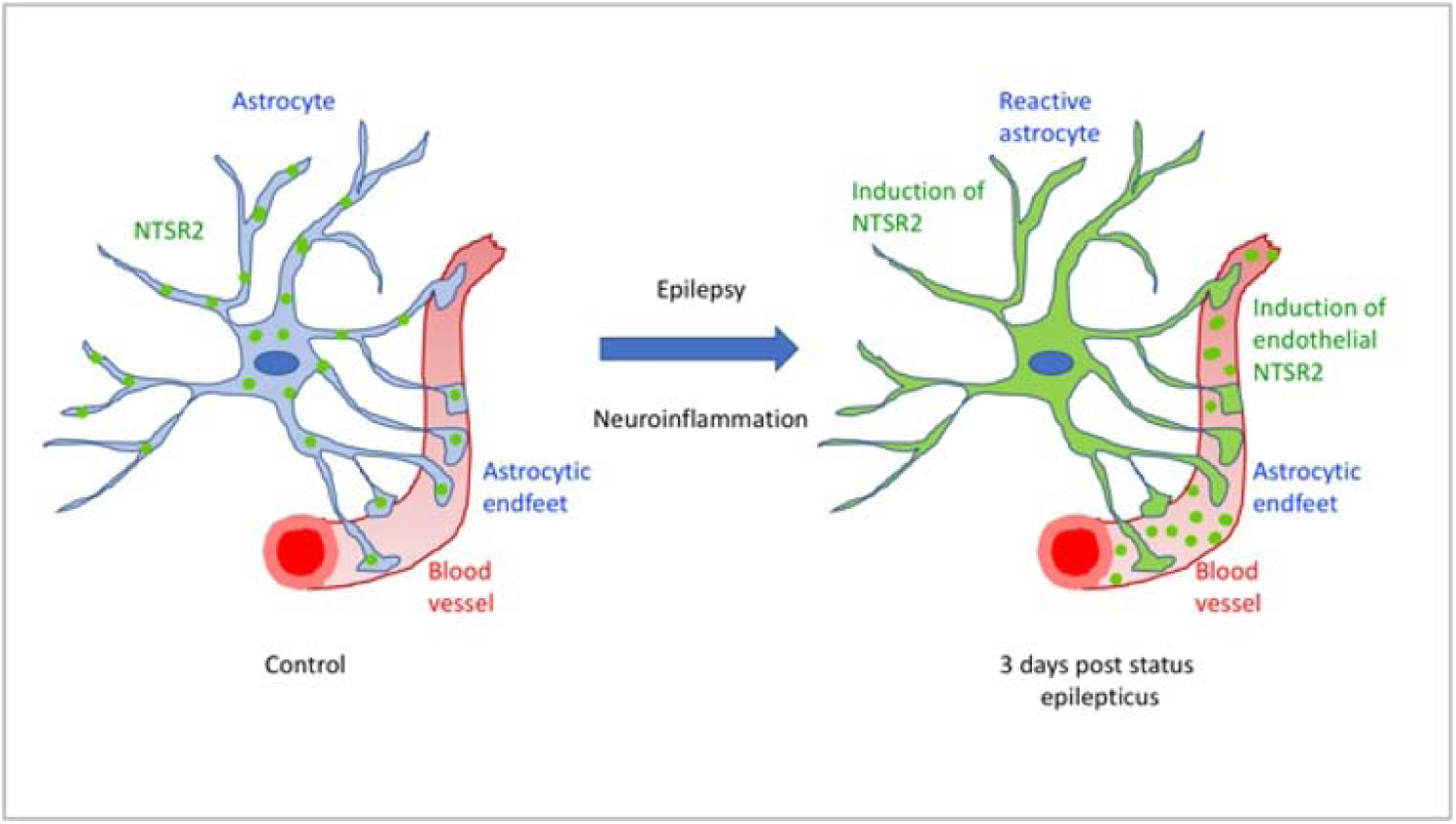

## INTRODUCTION

Neurotensin (NT) is a tridecapeptide highly conserved between species. In the periphery it is expressed in the gastrointestinal tract where it acts as a hormone, influencing food intake, body weight gain and the control of energy homeostasis. In the CNS, NT is a primary neurotransmitter/neuromodulator that mediates analgesia and is implicated in memory consolidation and in learning processes (Goedert and Emson, 1983; Vincent, 1995; Friry et al., 2002; Devader et al., 2006; Yamauchi et al., 2007). NT has been involved in a number of psychiatric pathologies, distinctively schizophrenia (Sharma et al., 1997), eating disorders (Hawkins et al., 1986), Parkinson’s disease (Hernandez-Chan et al., 2015), glioma (Ouyang et al., 2015), pain (Williams et al., 1995), cerebral ischemia (Torup et al., 2003) and epilepsy (Lee et al., 2009; St-Gelais et al., 2006). Previous studies have interrelated NT and seizures. Indeed, in the KA model of epilepsy, NT-like immunoreactivity in the rat hippocampus was reduced after onset of seizures, suggesting NT release (Sperk et al., 1986). Intracerebroventricular (i.c) or intraperitoneal (i.p) administration of NT or its analogues showed anti-convulsant effects in a corneal stimulation mouse model of pharmaco-resistant epilepsy (Lee et al., 2009; Green et al., 2010). However, the mechanism by which NT receptors (NTSRs) mediate anti-convulsive effects remains unknown. NT and its analogues induce generalized hypothermia of 2-4°C when administered directly in the CNS by intracisternal or i.c administration (Nemeroff et al., 1980). This reduction in body temperature prevents hippocampal neuronal damage and preserves locomotor activity during an ischemic insult, highlighting NT’s therapeutic potential (Babcock et al., 1993).

Recently, we showed that intravenous (i.v) administration of NT analogues vectorized with a brain shuttle peptide targeting the blood brain barrier LDL receptor induced potent neuroprotective and anti-neuroinflammatory effects in the hippocampus of a rodent model of epilepsy. These neuroprotective effects of NT on hippocampal neurons were probably mediated by hypothermia, *in vivo*, but we demonstrated they could also be observed *in vitro*, independently of hypothermia (Soussi et al., manuscript in preparation).

NT mediates its central and peripheral effects by interacting with three receptor subtypes, namely NTSR1, NTSR2 and NTSR3. NTSR1 and NTSR2 are G protein-coupled receptors with seven transmembrane domains (Vincent, 1995). NTSR3 or gp95/Sortilin 1 is a type I receptor with a single transmembrane domain that belongs to the Vps10p-containing domain receptor family and is not coupled to a G protein (Mazella, 2001). To date, not much is known on the potential role of NT in modulating neuroprotection and neuroinflammation and which receptors are involved, in which brain structures and cell types. NTSR1 and NTSR2 show high and low affinity for NT, respectively (Tanaka et al., 1990; Chalon et al., 1996), and these two receptors are found in different cell types of the nervous system at different developmental stages. NTSR1 is expressed prenatally in many rat brain structures. Expression peaks shortly after birth and is decreased in adulthood (Palacios et al., 1988). On the other hand, NTSR2 expression is essentially postnatal and increases progressively with age (Sarret et al., 1998; Lépée-Lorgeoux et al., 1999), raising interest on the role of this receptor in brain physiopathology.

NTSR2 is a 45 kDa protein in humans and rodents, endowed with lower affinity for NT compared to NTSR1 (Kd = 3–10 nmol/L compared to 0.1–0.3 nmol/L for NTSR1) (St-Gelais et al., 2006). Besides its expression in pancreatic β cells (Béraud-Dufour et al., 2009) and human B lymphocytes (Saada et al., 2012), NTSR2 is essentially expressed in the CNS, including the hippocampus, cerebral cortex, cerebellum, olfactory bulb, substantia nigra, and ventral tegmental area (Walker et al., 1998; Lépée-Lorgeoux et al., 1999; Sarret et al., 2003). Globally, NTSR2 exerts the same effects as NTSR1 with a more characterized role in pain-reducing modulation (reviewed in Kleczkowska and Lipkowski, 2013) and fear memory (Yamauchi et al., 2007). Rat NTSR2 mRNA expression was shown in astrocytes *in vivo* (Walker et al., 1998; Yamauchi et al., 2007), and NTSR2 protein immunofluorescence was reported in astrocytes in the ventral tegmental area in mice (Woodworth et al., 2018). In contrast, other studies reported no NTSR2 immunostaining in astrocytes in adult rat brain (Sarret et al., 2003). Thus, the neurocellular expression of NTSR2 remains controversial, and up to now, the data available is limited.

In the present work, we assessed the expression of NTSR2 in the hippocampus, both in physiological and pathological conditions. In particular, we questioned whether expression of the NTSR2 receptor was modulated in conditions of neuroinflammation using a pathophysiological model of TLE induced by pilocarpine in adult rats. In this model, animals undergo a rapid and intense neuroinflammatory reaction with a pattern of reactive gliosis in the hippocampus involving the activation of microglia and astrocytes (Clifford et al., 1987; Garzillo & Mello, 2002).

Our study shows for the first time unambiguous expression of NTSR2 protein in hippocampal astrocytes of the adult rat, both *in vivo* and *in vitro*. In the pilocarpine epileptic rat model, NTSR2 expression was increased rapidly when neuroinflammation peaks, and decreased at the chronic phase (3 months), when astrocyte reactivity subsides (Garzillo and Mello, 2002; Choi and Koh, 2008). NTSR2 was found also to be expressed in microglial cells *in vitro* but not *in vivo*. We also report increased NTSR2 expression in blood vessels during inflammation and showed that a NT analogue that elicits anti-neuroinflammation effects *in vivo* downregulated glial reactivity *in vitro*. In all, our work demonstrates the involvement of the neurotensinergic system in parenchymal astrocytes and in the glio-vascular unit during neuroinflammation and suggest that targeting the NTSR2 receptor could modulate neuroinflammation.

## MATERIALS AND METHODS

### Experimental Animals

All experimental procedures involving rats and mice were approved by National and European regulations (EU directive N° 2010/63) and in agreement with the authorization for animal experimentation attributed to the laboratory by the Prefecture des Bouches du Rhône (permit number: D 13 055 08) and to the project (N° 00757.02) by the French Ministry of Research and Local Ethics Committee. Animals were maintained in the animal facility with 12H light-dark cycles, had access to food and water *ad libitum* and were treated according to Appendix A of the European Convention for the Protection of Vertebrate Animals used for Experimental and other Scientific Purposes, ETS No. 123. All efforts were made to minimize animal suffering and to reduce the number of animals used.

### Rat pilocarpine model

Adult male Wistar rats weighing 200-290 g (Charles River, France) were first injected i.p. with a low dose of the cholinergic antagonist scopolamine methyl nitrate (2 mg/kg; Sigma, Saint Louis, MO, USA), in order to minimize the peripheral effects of pilocarpine hydrochloride (320 mg/kg; Sigma), a muscarinic cholinergic agonist that we diluted in 0,9% NaCl and administered i.p.30 min after scopolamine methyl nitrate. Control rats received an injection of 0,9% NaCl. The injection protocols were similar to those previously described for the generation of TLE (Mello et al., 1993; Obenaus et al., 1993; Dinocourt et al., 2003; Sbai et al., 2012; Soussi et al., 2015). Only animals that developed confirmed SE after the pilocarpine injection were included in the study. To reduce animal mortality, SE was stopped after 1H by two injections of Diazepam (Valium) with a 15 min interval (10 mg/kg, i.p. Roche, France). Pilocarpine-treated animals were then observed periodically for general behavior and occurrence of spontaneous seizures. Pilocarpine-treated animals were studied at several post-injection intervals: during the latent period, when animals displayed an apparently normal behavior (3 days, 1 and 2 weeks, n ≥ 6 at each time point), and during the chronic stage, when the animals developed spontaneous recurrent limbic seizures (12 weeks; n ≥ 6). During the following days after induction of SE, animals were nurtured and assisted to drink water. For the weakest ones, a double dose of soluble NaCl 0,9 % and glucose (10 mg/kg, i.p.) was provided twice per day until they regained their weight or reached the endpoint. Animals were housed two per cage in enriched environment to minimize stress prior to SE and one per cage to avoid aggressive behaviors post-SE.

### Mouse kainic acid model

As an alternative model of epilepsy, young adult male FVB/N mice (Janvier Laboratories, France), 25-30 g, 9-10 weeks old, were injected subcutaneously with a single dose of kainic acid (KA, 40-45 mg/kg; Abcam, France) to generate mice with spontaneous recurrent seizures as a hallmark of SE as previously described (Schauwecker & Steward, 1997). All animals were housed six per cage. KA-injected mice were individually housed and received a 0,5 ml i.p. dose of glucose and also had free access to Doliprane (paracetamol, Sanofi, Gentilly, France) at 2,4 % in agarose gel in order to reduce pain. Mice were observed during 9H for onset and extent of seizure activity and only mice developing typical SE were included in the study. Body weight was monitored daily.

### Tissue Preparation

Rats were deeply anesthetized with pentobarbital sodium injection (Nembutal, 120 mg/kg, i.p., Ceva, France) and perfused through the heart with a fixative paraformaldehyde-based solution (Antigenfix, Diapath, Italy). The brains were then removed from the skull 30min later, and were post-fixed in the same fixative solution for 24H at 4°C. Finally, brains were rinsed 3x in Phosphate Buffer (0,12 M; PB, pH 7,2-7,4) and cryoprotected in 30% sucrose solution in PB 0,12 M until fully dehydrated (indicated by sinking of the brain at the bottom of the pot, lasting from 2-4 days). Brains were first frozen in isopentane solution (Sigma) at −80°C and then fixed in O.C.T. Tissue Tek (Sakura Finetek, Torrance, CA, USA) and sectioned coronally at 40 µm with a cryostat. The sections were collected sequentially in wells of culture plates containing an ethylene glycol-based cryoprotective solution and stored at −20°C until processing. Selected sections from each rat covering the dorsal hippocampus (from bregma −3,30 to −4,80 mm according to the rat brain Atlas (Paxinos & Watson, 1998), were used for immunohistochemical staining. Free-floating sections from control (CTL) and pilocarpine-treated rats (PILO) were always processed in parallel.

Mice were deeply anesthetized with chloral hydrate (700 mg/kg; ProLabo, France) and transcardially perfused with Antigenfix solution. The brains were extracted and post-fixed for 1H at room temperature (RT) and rinsed in PB 0.12 M. Forty μm coronal sections were generated using a vibratome, immersed in a cryoprotective solution and stored at −20°C until use for immunofluorescence. From each mouse, selected sections from the dorsal hippocampus (bregma −1,55 to −2,35 according to the mouse brain Atlas (Paxinos & Franklin, 2019)), were processed for immunohistochemistry. Mice were sacrificed at 7 days post-SE (KA 7D) and free-floating sections from CTL and KA-treated mice were processed in parallel.

### Immunohistochemistry

#### Double immunofluorescence labeling for NTSR2 and GFAP/Iba1

For double labeling of primary antibodies originating from the same host species, in our case rabbit, a protocol from Jackson Immunoresearch was used and optimized for our purposes. Free-floating sections from CTL and pilocarpine-treated rats (PILO rats) from the dorsal hippocampus were processed in parallel. References of primary antibodies used in this study are summarized in Table 1. Free-floating sections were permeabilized in a solution containing 3% Bovine Serum Albumin (BSA) and 0,3% Triton X-100 in PB 0,12 M for 1H at RT. Following permeabilization, sections were first incubated overnight at 4°C with rabbit anti-NTSR2 diluted in a blocking solution containing 3% BSA diluted in PB 0,12 M. The next day, slices were washed three times in PB 0,12 M under agitation and incubated with secondary antibody, goat anti-rabbit biotin (1:200) (Life Technologies, France) diluted in the blocking solution for 2H, and then revealed with streptavidin IgG conjugated with AlexaFluor 488 (Jackson Immunoresearch, West Grove, PA, USA) diluted in blocking solution (1:800) for 2H, in dark. Next, slices were washed three times in PB 0,12 M and incubated with normal rabbit serum (1:20) (Jackson Immunoresearch) diluted in PB 0,12 M for 2H at RT, in dark, to saturate open binding sites on the first secondary antibody with IgG. Following three washes in PB 0.12 M, slices were incubated with unconjugated Fab goat anti-rabbit IgG (H+L) (20 μg/mL) (Jackson Immunoresearch) diluted in PB 0,12 M for 2H at RT, in dark, to cloak the rabbit IgG and hamper binding of the second secondary antibody. Afterwards, slices were incubated overnight in the dark at 4°C under agitation with rabbit anti-GFAP or rabbit anti-Iba1. The last day, slices were washed three times in PB 0,12 M and incubated with a goat anti-rabbit IgG (H+L) highly cross-adsorbed AlexaFluor A594 (1:1.000, Life Technologies) diluted in blocking solution for 2H, in dark. Nuclei were counterstained with 5 μg/mL 4, 6- Diamidino-2-phenylindole (DAPI, Sigma) for 0,5H at RT, in dark. After three washes in PB 0,12 M, floating sections were mounted on Superfrost Plus glass slides using Fluoromount-G Mounting medium (Electron Microscopy Sciences, Hatfield, PA, USA) and stored at −20°C until imaging and analysis. The specimens were then analyzed using a confocal microscope (Zeiss, LSM 700, Jena, Germany) and images were acquired using Zen software (Zeiss), and processed using Adobe Photoshop and *ImageJ* softwares.

**TABLE 1.** Primary antibodies used for immunofluorescence analysis

| **Antibodies** | **Species** | **Sources and references** | **Dilution** |
| --- | --- | --- | --- |
| NTSR2 | Rabbit | Novus Biologicals, NBP1-00959 | 1:100 |
| GFAP | Rabbit | Dako, Z0334 | 1:500 |
| Iba1 | Rabbit | Wako, 019-19741 | 1:500 |
| PDGFR | Rabbit | Abcam, ab32570 | 1:100 |
| Collagen IV | Goat | Southern Biotech, 1340-01 | 1:250 |
| CD31 | Mouse | Abcam, ab64543 | 1:100 |
| MAP2 | Chicken | Abcam, ab5392 | 1:500 |

The specificity of the double immunohistochemical labeling for antibodies originating from the same host such as NTSR2 and GFAP or Iba1 was tested by incubating some sections from CTL and pilocarpine-treated animals with two primary antibodies with different targets; GFAP (rabbit, for astrocytes) and Iba1 (rabbit, for microglia). The two antibodies gave a clearly distinctive specific pattern and did not overlap. To judge the efficiency of the blocking, the second primary antibody was omitted, but not the corresponding secondary antibody. In all cases, the second secondary antibody did not detect the first primary antibody. Additional controls included incubation of some sections in a mixture of one primary antibody and biotinylated rabbit normal IgG, or rabbit normal IgG. The pattern of immunolabeling of these sections was the same as for sections processed for single labeling.

#### Triple immunofluorescence labeling for CollV, CD31, NTSR2

Following permeabilization and blocking, sections were incubated overnight at 4°C under agitation with goat anti-ColIV diluted in blocking solution. Next day, sections were washed three times in PB 0,12 M under agitation and then incubated overnight at 4°C with the other two primary antibodies, mouse anti-CD31 and rabbit anti-NTSR2, sequentially (i.e. one per day). The fourth day, sections were washed three times in PB 0,12 M under agitation and then incubated for 2H, in the dark with the appropriate secondary antibodies: donkey anti-goat IgG (H+L) highly cross-adsorbed AlexaFluor A488 for ColIV, donkey anti-mouse IgG (H+L) highly cross-adsorbed AlexaFluor A594 for CD31 and donkey anti-rabbit IgG (H+L) highly cross-adsorbed AlexaFluor A647 for NTSR2 (Life Technologies), diluted in blocking solution at 1:800. Slices were then washed three times in PB 0,12 M under agitation and nuclei were counterstained with 5 μg/mL DAPI for 0,5H at RT, in dark. Finally, sections were mounted using Fluoromount-G Mounting medium and stored at −20°C. The specificity of the triple immunohistochemical labeling was validated by incubating some sections in a solution omitting the primary antibodies in the presence of the appropriate secondary antibodies. No staining was detected under these conditions.

#### Triple immunofluorescence labeling PDGFRβ, CD31, NTSR2

For the immunostaining of PDGFRβ and NTSR2, sections were processed as for NTSR2 and GFAP/Iba1 immunostaining. Briefly, sections were incubated with rabbit anti-PDGFRβ and the next day after several rinses in PB 0,12 M, they were incubated sequentially with goat anti-rabbit biotin, streptavidin IgG conjugated with AlexaFluor A488 (Jackson Immunoresearch), normal rabbit serum, Fab goat anti-rabbit IgG, and finally incubated with rabbit anti-NTSR2 overnight at 4°C. The third day, following three washes in PB 0,12 M, slices were incubated with mouse anti-CD31 overnight. The last day, slices were washed three times and incubated with goat anti-mouse IgG1 (H+L) cross-adsorbed AlexaFluor A594 (1:800) for CD31, and goat anti-rabbit IgG (H+L) highly cross-adsorbed AlexaFluor A647 (1:1.000) (Life Technologies) in blocking solution for 2H, in dark. Nuclei were counterstained with DAPI and sections were mounted using Fluoromount-G Mounting medium and stored at −20°C. The specificity of the triple immunohistochemical labeling was tested as described above. In all cases, no specific staining was detected under these conditions.

#### Double immunofluorescence labeling for NTSR2 and MAP2

Sections were incubated overnight at 4°C under agitation with rabbit anti-NTSR2 diluted in blocking solution. Next day, they were washed three times in PB 0,12 M under agitation and incubated overnight at 4°C with chicken anti-MAP2. The last day, sections were washed 3 times in PB 0,12 M under agitation and then incubated for 2H, in the dark with the appropriate secondary antibodies: goat anti-rabbit IgG (H+L) highly cross-adsorbed AlexaFluor A488 for NTSR2 and goat anti-chicken IgY (H+L) AlexaFluor A594 (Life Technologies) for MAP2 diluted in blocking solution at 1:800. Counterstaining with DAPI, mounting, storage and imaging were as described above.

### Primary glial cell cultures

Glial cells (astrocytes, microglia) were prepared from embryonic day 18 (E18) or newborn (P0) rats cortices. Briefly, rats were killed by decapitation, brains were removed and cortices were dissected out and cleared from the meninges. Pieces of cortices were digested in a Trypsin-EDTA 1x solution for 15 min at 37°C and rinsed three times in HBSS 1X followed by three rinses in Dulbecco’s Modified Eagle Medium Glutamax 1X (DMEM) medium supplemented with 10% (v/v) heat-inactivated Fetal Bovine Serum (FBS). Next, cortices were mechanically dissociated by successive pipetting in DMEM supplemented with FBS. Dissociated cells were grown in DMEM supplemented with 10% FBS and 100 i.u./mL penicillin/streptomycin (all supplements were from Thermo Fisher Scientific, France), in a humidified atmosphere containing 5% CO_2_, at 37°C. The medium was changed twice per week and the cultures were used for experiments after 15 days in culture.

### Induction of inflammation and pharmacological treatments in primary glial cultures

Inflammation in primary glial cultures was induced by the addition of proinflammatory agents such as interleukin 1 beta (IL1β, 10 ng/mL; PeproTech, France) or lipopolysaccharide (LPS, 1 μg/ml; Sigma) for 1, 6, and 24H, at 37°C. Cultures were also incubated with 1 μM VH-N412 for 24H in the presence or absence of IL1β or LPS. For studies of immediate-early gene responses, primary glial cultures were incubated with the protein synthesis inhibitor, cycloheximide (CHX, 10 μg/ml, Cell Signaling Technology, MA, USA) in the presence or absence of LPS for 1, 6, and 24H at 37°C. Following treatments, after one wash with PB 0,12 M, cells were either fixed with 4% paraformaldehyde (PFA) in PB 0,12 M for 20 min at RT and then processed for immunocytochemistry, or used for RNA extraction and RT-qPCR analyses.

### Immunocytochemistry

Cell coated coverslips were rinsed three times with PB 0,12 M and permeabilized in a blocking solution containing 3% BSA and 0,1% Triton X-100 diluted in PB 0,12 M for 30 min at RT. Then, coverslips were incubated overnight with rabbit anti-NTSR2 antibody placed on droplets of 50 μl of antibody solutions diluted in the 3% BSA blocking solution with the cells side facing down, inside a humidity chamber at 4°C. The next day, coverslips were washed three times in PB 0,12 M under agitation at RT and incubated with goat anti-rabbit IgG (H+L) highly cross-adsorbed AlexaFluor A488 (1:800, Life Technologies) diluted in the blocking solution for 2H, in dark. After three washes in PB 0,12 M, coverslips were incubated with normal rabbit serum (1:20) for 2H and then with Fab goat anti-rabbit IgG (20 μg/mL) for another 2H at RT, in dark, before incubation with rabbit anti-GFAP or rabbit anti-Iba1 at 4°C overnight, in dark. The last day, coverslips were washed three times in PB 0,12 M and incubated with a goat anti-rabbit IgG (H+L) highly cross-adsorbed AlexaFluor A594 (1:1.000, Life Technologies) antibody diluted in the blocking solution for 2H at RT, in dark. Nuclei were counterstained with 5 μg/mL DAPI for 0,5H at RT, in dark. After three washes in PB 0,12 M, coverslips were rapidly rinsed three times in dH_2_O and let to dry in dark, before mounting on Superfrost glass slides using Fluoromount-G Mounting medium and stored at −20°C.

Labeling specificity was assessed under the same conditions, by incubating some coverslips from control and treated cells in a solution omitting one of the primary antibodies, and, furthermore, the specificity of the double immunolabeling for antibodies originating from the same host was evaluated by incubating some coverslips with two primary antibodies with different targets, GFAP and Iba1. In all cases, no overlap of antibodies was detected.

### MCP-1 (CCL2) assay by ELISA

Primary glial cell cultures were stimulated with IL1β (10 ng/ml) or LPS (1 μg/ml) for 24H. Then, supernatants were collected, centrifuged, and stored at −80°C until analysis. Monocyte chemoattractant protein 1/chemokine C-C motif ligand 2 (MCP1/CCL-2) levels were evaluated using a commercially available ELISA kit (PeproTech) according to the manufacturer’s instructions. All samples were analyzed in triplicate. The detection threshold was 16 pg/ml of cytokine.

### RNA Extractions and Taq Man quantitative PCR (qPCR)

Total RNA was prepared from primary rat glial cultures using the Nucleospin RNA plus kit (Macherey Nagel, France). cDNA was synthesized from 500 ng of total RNA using the High Capacity RNA-to-cDNA Kit (Applied Biosystems, CA, USA). For real-time qPCR, 12,5 ng of cDNA were used. The samples were run in duplicate on 96-well plates and then analyzed with 7500 v2.0 software (Applied Biosystems). The conditions of the thermal cycle were as follows: initial denaturation at 95°C for 40 Cycles, denaturation at 95°C, and hybridization and extension at 60°C. Relative expression levels were determined according to the ΔΔCt (Ct: cycle threshold) method where the expression level of the mRNA of interest is given by 2^ΔΔCT^, where ΔΔCT = ΔCt target mRNA − ΔCt reference mRNA (*GAPDH*, housekeeping gene) in the same sample. PCR experiments were performed with the 7500 Fast Real Time PCR System (Applied Biosystems), according to the manufacturer’s recommendations. All reactions were performed using TaqMan Fast Universal PCR Mix (Applied Biosystems) and TaqMan Assays (Applied Biosystems) probes (see Table 2).

**TABLE 2.** Rat TaqMan probes used for qPCR analysis

| **Gene name** | **Gene description** | **Probe ID** |
| --- | --- | --- |
| *Ntsr1* | Neurotensin receptor 1 | Rn01415846 |
| *Ntsr2* | Neurotensin receptor 2 | Rn00575514 |
| *Ntsr3/Sort1* | Neurotensin receptor 3/Sortilin 1 | Rn01521847 |
| *Mcp1/ccl2* | Monocyte Chemoattractant ligand 1 | Rn00580555 |
| *Il1b* | Interleukin 1 | Rn00580432 |
| *Gfap* | Glial Fibrillary Acidic Protein | Rn01253033 |
| *Zif-268/egr1* | Zinc finger protein 225/early growth response protein 1 | Rn00561138 |
| *Gapdh* | Glyceraldehyde-3-phosphate dehydrogenase | Rn01775763 |

### Synthesis of VH-N412

VH-N412 (Pr-[cMPipRLRSarC]_c_-PEG6-RRPYIL-OH) was synthesized by solid phase peptide synthesis method using standard Fmoc/tBu strategy and a Fmoc-Leu-Wang Resin on a Liberty™ (CEM) microwave synthesizer. In brief, resin-bound peptide was cleaved using a solution comprised of TFA/TIS/H_2_O/EDT: 94/2/2/2 for 2H at RT. Crude peptide was then precipitated using ice-cold ether, centrifuged at 3000 rpm for 8 min and lyophilized in H_2_O/0.1%TFA to obtain a white powder. Crude linear Pr-cMPipRLRSarC-PEG6-RRPYIL-OH conjugate was then dissolved in AcOH 0.5% to reach 0.5 mg/mL final concentration. Ammonium carbonate (2N) was added to the conjugate solution to reach an approximate basic pH of 8-9. K_3_[Fe(CN)_6_] (0.01N) was then added to the reaction mixture until a bright and persistent yellow color was observed to allow mild oxidation conditions. Monitoring of the disulfide bond formation between the two cysteines was performed by analytical RP-HPLC. After 30 min, the reaction mixture was filtered over 0.2µM and purified by preparative RP-HPLC. The pull of fractions containing >95% pure VH-N412 was freeze-dried to yield a pure white to yellow powder (purity > 95% assessed by analytical RP-HPLC) and MALDI-TOF (m/z) for C94H163N27O24S3, [M+H]+ calc. 2151.15, found 2151,11 Da.

### Data analysis

All acquisitions were performed using a confocal laser-scanning microscope (LSM 700, Zeiss) through a x20 or x40 oil objective and analysis of immunostaining images was performed using ZEN software (Zeiss). To visualize the whole hippocampus, the mosaic function was required. Z-stack function was also useful to determine precisely the co-expression of two markers in the same cells. Finally, *ImageJ* software (NIH) was used to quantify each immunolabeling. Pictures were binarized to 16-bit black and white images and a fixed intensity threshold was applied, defining each staining. Images were obtained with double averaging, frame size 1024×1024, and 1 AU pinhole for each channel.

#### Quantification of NTSR2 levels in GFAP-positive (GFAP+) cells on tissue

The mean fluorescence intensity (mean grey value) was measured in three different brain regions: dentate gyrus (DG), covering the hilus (H), granule cell layer (G), and the inner- and outer molecular layer (IML, OML), CA1, and CA3. Cells were identified as GFAP+ by an apparent nucleus-surrounding structure by co-labeling of NTSR2 with GFAP, and measurement of fluorescence was made in Z-stack x40 acquired images on the stack in which the intensity of NTSR2 was the highest. Measurement of fluorescence was made on the cell body and processes, when these were well defined, by drawing around them. A minimum of 200 GFAP+ cells were analyzed from three pilocarpine-treated animals at each time point and three CTL (2 sections/animal) after subtracting the background fluorescence. Background fluorescence was set in areas devoid of stained cells in the same sections an average value of 3 such areas was obtained from every image. Data are presented as the mean percentage of fluorescence of NTSR2+ cells co-labeled with GFAP normalized to the controls [mean ± SEM (standard error of the mean)].

#### Quantification of NTSR2 levels in blood vessels on tissue

The mean fluorescence intensity was measured in blood vessels of the DG and the lacunosum-moleculare (LM), by drawing and measuring around the inner and outer surface of each vessel. Measurement of fluorescence was made in Z-stack x20 mosaic images on the stack in which the intensity of NTSR2 was the highest. All vessels from three animals from CTL and PILO 3D were quantified (55 and 79 vessels, respectively). Background fluorescence was subtracted. Background fluorescence was set in the stratum radiatum, in areas devoid of brain vessels in the same sections an average value of three such areas was obtained from every image. Data are presented as the mean percentage of fluorescence of NTSR2+ cells co-labeled with GFAP normalized to the controls (mean ± SEM).

#### Quantification of NTSR2, GFAP and Iba1 levels in CTL, inflamed and VH-N412-treated primary glial cultures

The mean fluorescence intensity of NTSR2, GFAP and Iba1 in the cell body and processes was measured in double stained cells for NTSR2 and either GFAP or Iba1. Measurement of fluorescence was made in Z-stack x20 acquired images on the stack where the intensity of each marker was the highest, on isolated cells. A minimum of 100 GFAP+/Iba1+ cells were analyzed for each condition from a total of 10 images/condition spanning the whole coverslip, randomly acquired, from three independent experiments, after subtracting the background fluorescence. Background fluorescence was set in areas devoid of cells in every image. Data are presented as the mean percentage of fluorescence of NTSR2 or GFAP/Iba1 in the same cells, normalized to the controls (mean ± SEM).

### Statistical analysis

All experiments were performed at least three times with different rat series or independent cultures. Student’s *t*-test was used to compare two groups. The quantification of mean values and double-positive cells was analyzed with a one-way ANOVA, followed by Tukey’s *post hoc* test for multiple comparisons. All data are expressed as the mean ± SEM. Statistical significance was set up to * *p*<0,05, ** *p*<0,01, and *** *p*<0,001.

## RESULTS

### NTSR2 protein is expressed in the rat hippocampus and increases following pilocarpine-induced SE

Several lines of evidence suggest that NTSR2 mRNA is expressed in glial cells, namely astrocytes, *in vitro* and *in vivo,* including in the hippocampus (Walker et al., 1998; Lépée-Lorgeoux et al., 1999; Nouel et al., 1999). The NTSR2 protein was not expressed in hippocampal astrocytes (Sarret et al., 2003) but was recently described in astrocytes of the ventral tegmental area (Woodworth et al., 2018). We questioned the expression of NTSR2 protein in the hippocampus of CTL rats and its modulation in reactive astrocytes associated with epilepsy and neuroinflammation. In particular, we studied NTSR2 expression pattern and distribution in the hippocampus of rats at different stages following pilocarpine administration. Indeed, this model is characterized by rapid astrocyte and microglial activation in the hippocampus (Shapiro et al., 2008).

All rats that were injected with pilocarpine and survived developed SE (mortality rate was around 25%). From 10 minutes to 1H following pilocarpine injection, rats exhibited limbic motor seizures every few minutes. After three weeks, spontaneous recurrent seizures started to appear that could last up to 60 sec; these developed into generalized seizures within the following days that persisted for the lifetime of the animals. These observations are in agreement with previous studies (Goffin et al., 2007; A. K. Sharma et al., 2007; Drislane et al., 2009; Knake et al., 2009).

We performed dual immunohistochemical labeling of NTSR2 (green) and glial fibrillary acidic protein (GFAP, red), a mature astrocyte marker (Dusart et al., 1991), at different stages following SE in pilocarpine-treated rats (Fig. 1). We chose to illustrate the results obtained at three days post SE (PILO 3D) with NTSR2 (Figs. 1 A,D) and GFAP (Figs. 1 B,E) labeling in control (CTL) (Figs. 1 A-C) and pilocarpine-treated rats (Figs. 1 D-F). Indeed, in our conditions, the three -day time point corresponds to strong astroglial inflammation, characterized by a dramatic increase of GFAP, in agreement with the temporal profile described by Shapiro et al. (2008). In sections from CTL rats, and at low magnification, NTSR2 immunolabeling was fairly homogeneous in all areas and layers of the hippocampus (Figs. 1A,C) and showed a regional- and laminar-specific pattern within the hippocampus. Modest to moderate NTSR2 immunolabeling was found within all layers, including the stratum oriens (O), stratum radiatum (R) and the stratum lacunosum-moleculare (LM) of the CA1-CA2-CA3 areas, the stratum lucidum (SL) of CA2-CA3, the molecular layer (M) and the hilus (H) of the dentate gyrus (DG) (Fig. 1A). The cell bodies of hippocampal pyramidal neurons (P) of CA1, CA2 and CA3, and granule cells (G) of DG were moderately stained. The NTSR2 immunolabeling pattern was clearly altered in PILO 3D compared to CTL rats. At low magnification, NTSR2 immunolabeling was characterized by a punctiform pattern in O, R, LM and H. Strong NTSR2 immunolabeling was observed within all layers and areas, including the O, R and the LM of the CA1-CA2-CA3 areas, the SL of CA2-CA3, the M, the cell bodies of P of CA1-CA2-CA3 areas, and the H of the DG (Figs. 1D,F).

**Figure 1.**
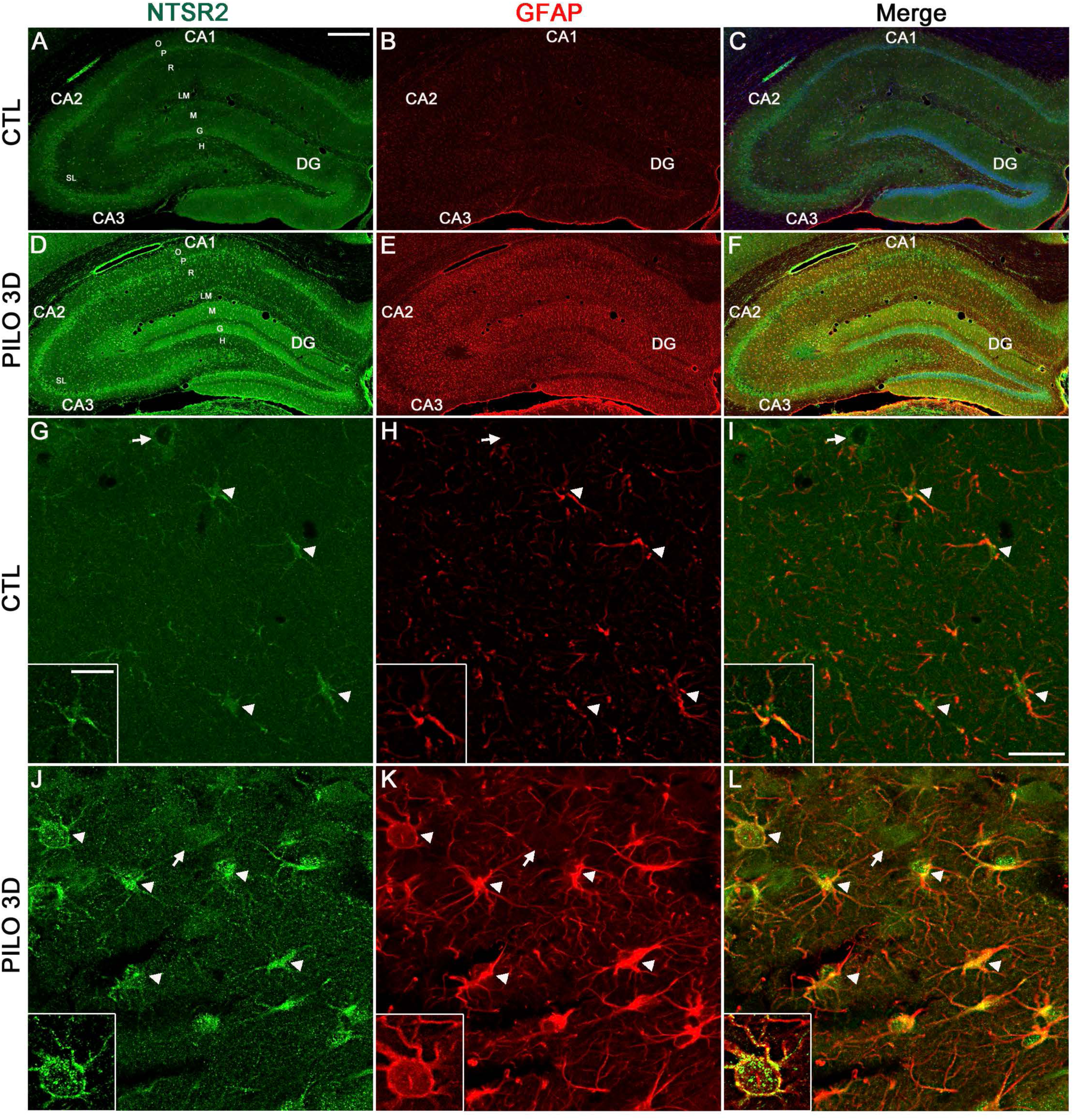
NTSR2 protein is mainly expressed in astrocytes and increases in all hippocampal areas following pilocarpine-induced SE. NTSR2 (green) and GFAP (red) double immunolabeling was performed in CTL (A-C, G-I) and pilocarpine-treated rats at day 3 post-SE (PILO 3D, D-F, J-L). Cell nuclei were counterstained with DAPI (blue, C and F). Merge corresponds to NTSR2/GFAP/DAPI (C and F). High magnification of the hilus of CTL (G-I) and PILO 3D (J-L) clearly shows NTSR2 immunolabeling in GFAP-positive astrocytes (see arrowheads), with stronger NTSR2 immunolabeling in PILO 3D compared to CTL animals. Note that non-astrocytic (GFAP-negative) NTSR2+ cells, most likely neuronal cells, are visible both in CTL and PILO 3D (G-L, see arrows). NTSR2 immunoreactivity is visible in the cell membrane, in the cell body and processes of astrocytes (see insets). In PILO 3D rats, NTSR2 immunolabeling displayed a more punctate pattern at the cell membrane as well as in the processes (Inset in J and L) compared to CTL rats. Panels I and L correspond to NTSR2/GFAP merged labelling. Scale bars: 450 μm in A-F, 10 μm in G-L and 5 μm in insets. O: stratum oriens; P: pyramidal neurons of CA1, CA2 and CA3; R: stratum radiatum; LM: stratum lacunosum-moleculare; M: molecular layer; DG: dentate gyrus; G granule cell layer of DG; H: hilus of the DG.

### NTSR2 protein is expressed in astrocytes and increases following pilocarpine-induced SE

At high magnification, all scattered GFAP positive cells located in the H were clearly immunolabeled for NTSR2, both in CTL and PILO 3D rats. GFAP immunolabeling was clearly enhanced in astrocytes of the DG, CA1, CA3, and the H of PILO 3D as compared to CTL rats, characteristic of astrocytic reactivity and neuroinflammation. NTSR2 was expressed in the cell bodies as well as within processes of hilar astrocytes of CTL and PILO 3D rats. While NTSR2 immunolabeling was fairly diffuse in CTL astrocytes (Figs. 1G-I, see insets), it appeared more punctiform in the astrocytes of the PILO 3D rats (Figs. 1J-L, see insets). NTSR2 staining was localized in part at the plasma membrane of astrocytes, both in CTL and PILO 3D rats, consistent with the cell membrane localization of NTSR2. In addition, NTSR2 immunolabeling of cell membranes as well as processes appeared markedly more intense in PILO 3D astrocytes compared to CTL astrocytes (compare insets in J and L with insets in G and I). All astrocytes, both in CTL and PILO 3D rats, as well as at all time points post-SE (data not shown), were NTSR2 positive.

Since differences in NTSR2 and GFAP labeling intensity in astrocytes were observed consistently between CTL and PILO 3D, we conducted semi-quantitative analysis at various time points post-SE in the DG, CA1 and CA3 to determine the relative extent of these changes (Fig. 2). In the DG, the mean NTSR2 levels were significantly increased in PILO 3D (129,7 ± 3,5%, 30%) and PILO 7D (119,1 ± 4,1%, 19%) when compared with CTL rats (100 ± 4,9%; n=3; *p*<0,01; Tukey’s test), whereas no difference was found in PILO 14D (105,1 ± 4,1%; n=3; *p*>0,05; Anova). The mean NTSR2 levels were significantly decreased at three months post SE (PILO 3M) (46,7 ± 2,5%, 53%; n=3; *p*<0,01; Tukey’s test) (Fig. 2A). In CA1, the mean NTSR2 levels in astrocytes were significantly increased in PILO 3D (182,3 ± 8,7%, 82%; n=3; *p*<0,01; Tukey’s test), PILO 7D (150,4 ± 5%, 50%) and to a much higher extent in PILO 14D (229,2 ± 8,9%, 129%) as compared to CTL (100 ± 4,9%; n=3; *p*<0,01; Tukey’s test). No difference was found in PILO 3M (79,3 ± 3,2%; n=3; *p*>0,05; Anova) (Fig. 2B). In CA3, the mean NTSR2 levels in astrocytes were significantly increased in PILO 3D (135,9 ± 4,1%, 36%), PILO 7D (128,2 ± 5,6%, 28%) and PILO 14D (123 ± 4,5%, 23%) as compared to CTL (100 ± 4,3%; n=3; *p*<0,01; Tukey’s test), whereas the mean NTSR2 levels in astrocytes were significantly decreased in PILO 3M (55 ± 2,8%, 45%; n=3; *p*<0,01; Tukey’s test) (Fig. 2C).

**Figure 2.**
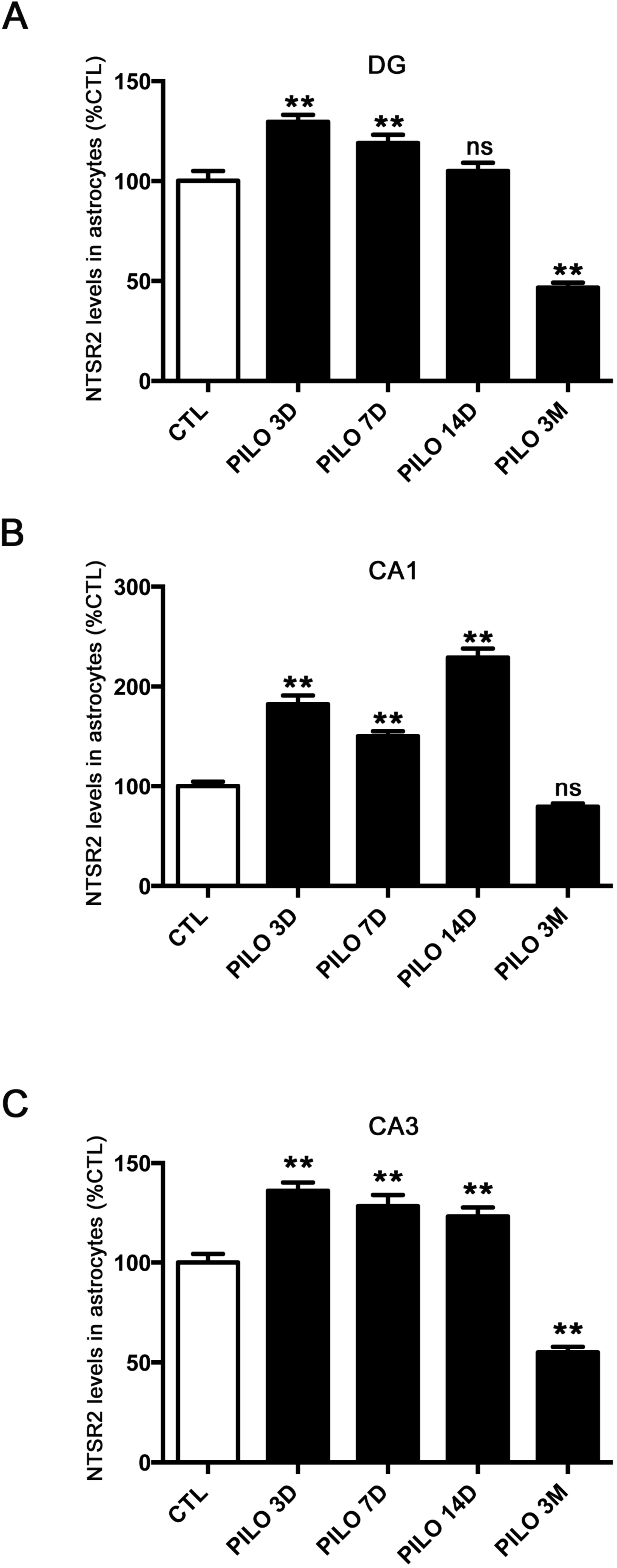
Mean intensities of NTSR2 immunolabeling in hippocampal astrocytes at different times following pilocarpine-induced SE, compared to CTL. In the DG (A) and CA3 (C), the highest increase for NTSR2 levels was at 3D post-SE, and at 14D for CA1 (B) compared to CTL. In all three areas, NTSR2 levels were higher in PILO vs CTL rats at all time points post-SE from 3 to 14D, except at 14D post-SE for DG. At 3 months post-SE, NTSR2 levels were significantly reduced in the DG and CA3 compared to CTL, but not in CA1. Astrocytes were detected by GFAP immunolabeling. NTSR2 levels were quantified in cells labeled with both NTSR2 and GFAP and values are given as the mean ± SEM as a percentage of CTL for each area. Asterisks indicate statistically significant differences: ** *p<*0,01 (one-way ANOVA followed by Tukey’s *post hoc* test).

### NTSR2 protein is expressed in the mouse hippocampus and increases following kainate-induced SE

In order to evaluate whether increased NTSR2 expression was specific to the rat pilocarpine model of epilepsy or may occur in other models, we injected KA to adult mice, generating SE and subsequent damage to the ipsilateral CA3 subfield, along with injury to the CA1 and DG/hilar region (Mouri et al., 2008). All mice developed spontaneous recurrent seizures within 5 days, in agreement with previously published studies (Mouri et al., 2008) and displayed sustained neuroinflammation with astrogliosis from 1 day to 1 month post-KA-induced SE (M. Ding et al., 2000; Z. Chen et al., 2005).

We performed dual immunohistochemical labeling of NTSR2 (green) and microtubule-associated protein 2 (MAP2, red), a mature neuronal marker (Izant & McIntosh, 1980), in CTL and KA-injected mice 1 week post-SE (KA 7D), when animals displayed sustained neuroinflammation (Fig. S1). In CTL mice, and at low magnification, NTSR2 immunolabeling was fairly homogeneous in all areas and layers of the hippocampus (Figs. S1A,C) and showed a regional- and laminar-specific pattern within the hippocampus. Modest to moderate NTSR2 immunolabeling was found within all layers, including the O, R and the LM of the CA1-CA2-CA3 areas, the SL of CA2-CA3, the M and the H of the DG (Fig. S1A). The cell bodies of P of CA1, CA2 and CA3, and G of DG were moderately stained. The NTSR2 immunolabeling pattern was clearly altered in KA 7D compared to CTL mice (Fig. 1SD). At low magnification, NTSR2 immunolabeling was characterized by a punctiform and filamentous pattern in O, R, LM and H. Strong NTSR2 immunolabeling was observed within all layers and areas, including the O, R and the LM of the CA1-CA2-CA3 areas, the SL of CA2-CA3, the M, the cell bodies of P of CA1-CA2-CA3 areas, and the H of the DG (Figs. S1D,F). This strong increase of NTSR2 is associated with remarkable decrease of MAP2 immunolabeling (compare Fig. S1B with Fig. S1E). At high magnification, all scattered NTSR2 positive cells located in the H were clearly immunolabeled for MAP2 in CTL mice. Such NTSR2 positive cells were not observed in KA 7D mice. Indeed, MAP2 immunolabeling was clearly decreased in neurons of the DG, CA1, CA2, CA3, and the H of KA 7D (Figs. 1SE,K) as compared to that of CTL mice (Figs. 1SB,H). This loss of MAP2 immunolabeling may be due to the neurodegenerative processes known to occur in this model (Mouri et al., 2008). However, few scattered neurons were observed within the hippocampal lesion sites, where neuronal loss was extensive. NTSR2 was expressed in the cell bodies as well as within the proximal dendrites of hilar neurons of CTL mice and in non-neuronal, MAP2-negative cells in KA 7D mice. At high magnification, NTSR2-positive cells had an intense fibrillose appearance with strong staining in the cell bodies and processes. These NTSR2-labeled MAP2-negative cells (Figs. S1J-L) were most likely astrocytes.

We next sought to determine whether NTSR2 is also expressed and/or regulated in microglial cells, that also contribute to inflammation in pilocarpine-induced seizures in rats. To this end, we performed double immunolabeling of NTSR2 (Figs. 3 A,D, green) and the ionized macrophage and microglia specific calcium-binding adaptor molecule-1 (Iba1) (Ohsawa et al., 2004) (Figs. 3B,E, red) in CTL and pilocarpine-treated rats at different time points post-SE, in the DG, CA1 and CA3. Similar to GFAP, increased Iba1 immunolabeling, characteristic of inflammation, was observed in hippocampal microglial cells of PILO 3D rats as compared to CTL rats (Shapiro et al., 2008). High magnification images from CTL rats (Figs. 3G-I) and PILO 3D (Figs. 3J-L), clearly showed that the NTSR2 and Iba1 antibodies stained distinct cells in the H of the DG (Figs. 3G-L, see arrowheads), and in all other subfields of the hippocampus (data not shown), suggesting that NTSR2 is not expressed in microglial cells *in vivo*. Altogether, our data show that NTSR2 protein is expressed in hippocampal astrocytes of CTL rats and its expression is up-regulated during inflammation in pilocarpine-induced seizures.

**Figure 3.**
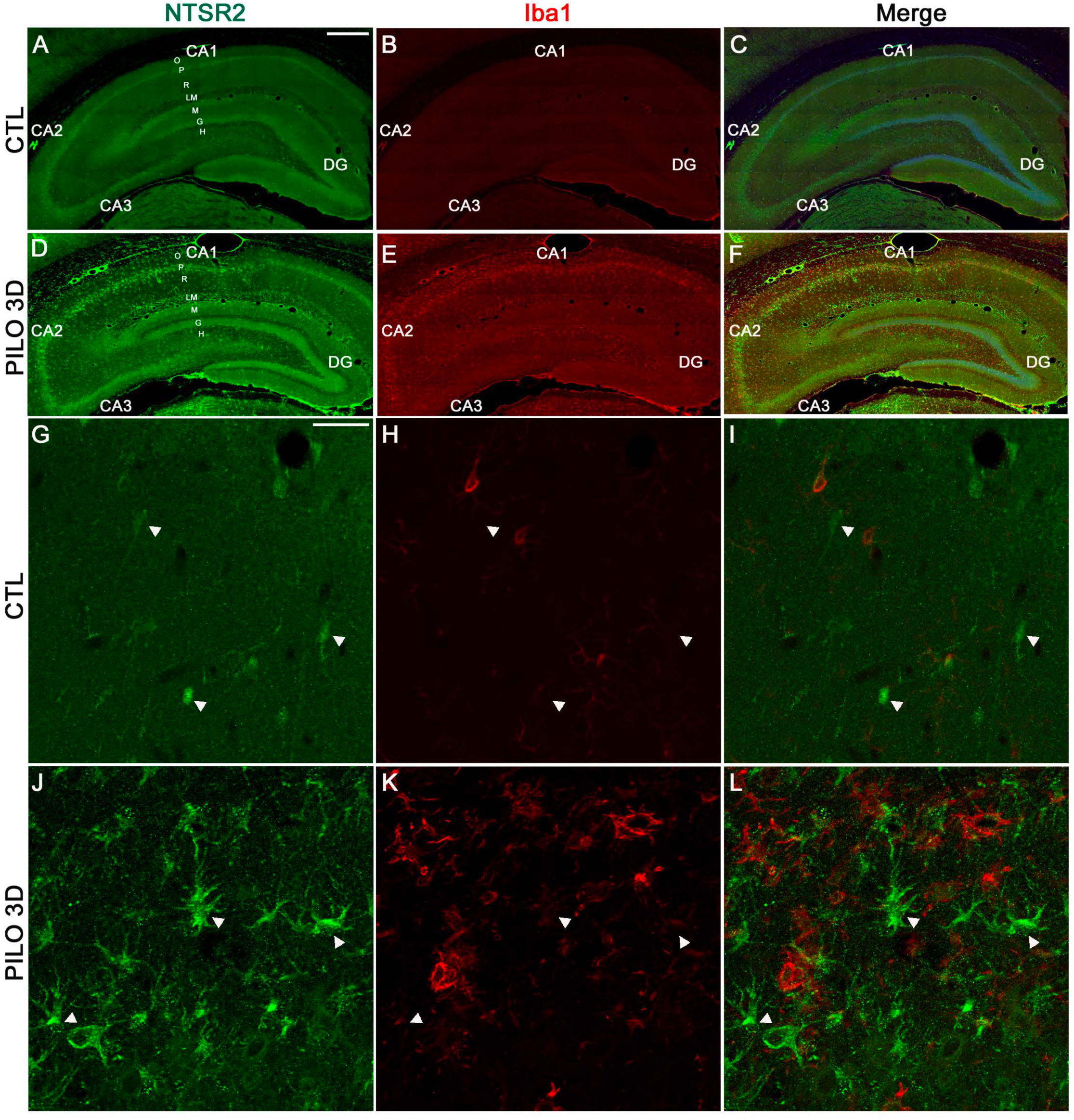
NTSR2 is not expressed in hippocampal microglial cells in CTL rats and following pilocarpine-induced SE. NTSR2 (green) and Iba1 (red) immunostaining was performed in CTL (A-C, G-I) and PILO 3D rats (D-F, J-L). Microglia were detected by Iba1 immunolabeling. Cell nuclei were counterstained with DAPI (blue, C and F). Merge corresponds to NTSR2/Iba1/DAPI (C and F). NTSR2 was not expressed in microglial cells since NTSR2 and Iba1 antibodies stain distinct cells as revealed by high magnification images (G-L, see arrowheads). Scale bars: 450 μm in A-F and 10 μm in G-L.

### NTSR2 protein is expressed in blood vessels and increases following pilocarpine-induced SE

Immunohistochemical analysis of hippocampal slices revealed that the NTSR2 protein is expressed in blood vessels of the LM of CTL (Fig. 4B, in green) and PILO 3D rats (Fig. 4E, in green). The pattern of NTSR2 immunolabeling was clearly altered in and around the blood vessels of PILO 3D rats (Figs. 4E,F), compared to that observed in CTL rats (Fig. 4B,C). The expression of NTSR2 increased in the blood vessels of the LM of PILO 3D. Furthermore, we observed several cells in the close vicinity of these vessels (see arrows), and other cells sparsely distributed around the blood vessels (see arrowheads), which displayed also high NTSR2 immunolabeling with a punctate pattern. Because differences in NTSR2 labeling intensity in the blood vessels were observed consistently between CTL and pilocarpine-treated animals, we conducted semi-quantitative analysis in CTL and PILO 3D rats to determine the relative extent of these changes (Fig. 4G). In the LM and DG, the mean intensities of labeling for NTSR2 were significantly increased in the PILO 3D animals (313,6 ± 21,4%, 213%; p<0,001, Student’s t-test), when compared with CTL rats (100 ± 12,2%). NTSR2-immunohistochemical labeling was abolished when the anti-NTSR2 antibody was omitted (data not shown). Thus, these data indicate upregulation of NTSR2 protein expression in the blood vessels during pilocarpine-induced seizures.

**Figure 4.**
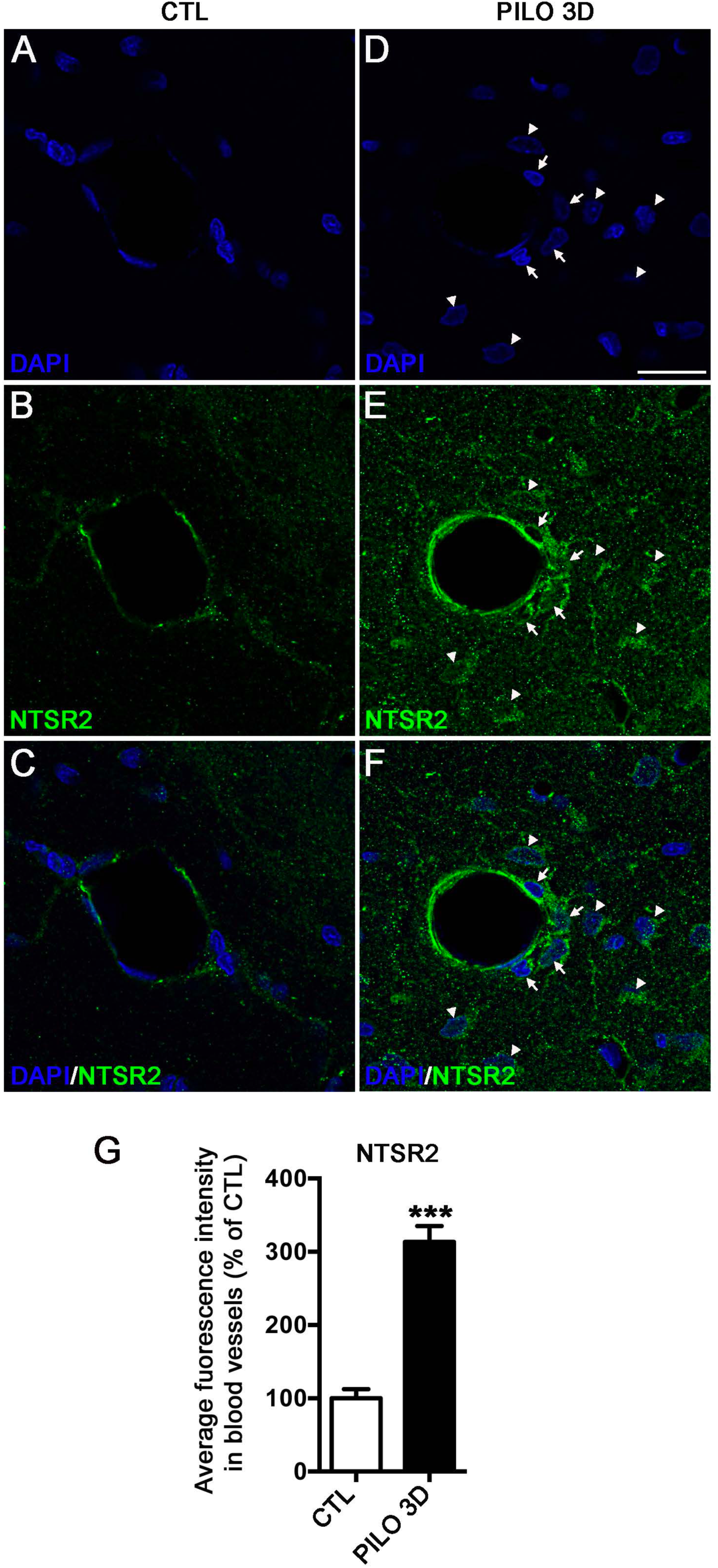
NTSR2 protein is expressed in blood vessels and increases following pilocarpine-induced SE. NTSR2 (green) immunostaining was performed in CTL (A-C) and PILO 3D rats (D-F). NTSR2 immunoreactivity was present in blood vessel cells of the LM with increased labeling in PILO 3D rats. In addition, several cells around these vessels displayed strong NTSR2 expression compared to CTL rats (arrows). Other dispersed cells in LM expressing NTSR2 are indicated (arrowheads). Cell nuclei were counterstained with DAPI (blue, A-F). Scale bar: 20 μm in all panels. G: Mean percentage of NTSR2 fluorescence intensity in DG and LM vessels in CTL and PILO 3D rats. Blood vessels of PILO 3D rats expressed significant NTSR2 levels compared to CTL rats vessels. Values are given as the mean ± SEM as a percentage of CTL. Asterisks indicate statistically significant differences: *** *p<*0.001 (Student’s *t*-test).

As for pilocarpine-treated rats, an increase of NTSR2 immunolabeling was also observed in the blood vessels of KA-treated mice (Fig. S2). Whereas in CTL mice the staining of NTSR2 around vessels was diffusively faint (Fig. S2B), in KA 7D mice NTSR2 immunoreactivity was strongly increased in the blood vessel cells of the LM compared to CTL mice (Fig. S2E). Several other cells around these vessels displayed differential, ranging from moderate to strong, NTSR2 expression, compared to CTL mice (arrowheads), while dispersed cells, likely perivascular astrocytes, projecting their processes (indicated by arrows and asterisks) were also observed around the blood vessels (Fig. S2F).

### NTSR2 protein is expressed in astrocytic end-feet and endothelial cells in blood vessels following pilocarpine-induced SE

Since all GFAP-labeled cells were also labeled for NTSR2, we performed high magnification confocal microscopic analysis to reveal the cellular compartments of astrocytes that are immunolabeled for NTSR2 (Fig. 5). We used GFAP to label astrocytes, astrocytic processes, and end-feet. Double immunofluorescence showed high NTSR2 expression in all reactive hypertrophic astrocytes (expressing high levels of GFAP) that were located around the blood vessels of the LM (Figs. 5 A-E, yellow, see arrows) in PILO 3D rats, when NTSR2 expression reached its peak. In addition, high magnification images revealed that astrocytic processes as well as their end feet that project on blood vessels were also strongly immunolabeled for NTSR2 (Figs. 5F-I, yellow, see arrowheads). Nevertheless, other cells that constitute the blood vessels that were not GFAP-positive also expressed high levels of NTSR2 (see stars in Figs. 5F-I). These vessels are constituted by specialized endothelial cells, surrounded by a basal lamina, pericytes, astrocyte-end feet that collectively form the blood-brain barrier (BBB), microglia and neurons (ElAli et al., 2014; Uzunalli et al., 2019).

**Figure 5.**
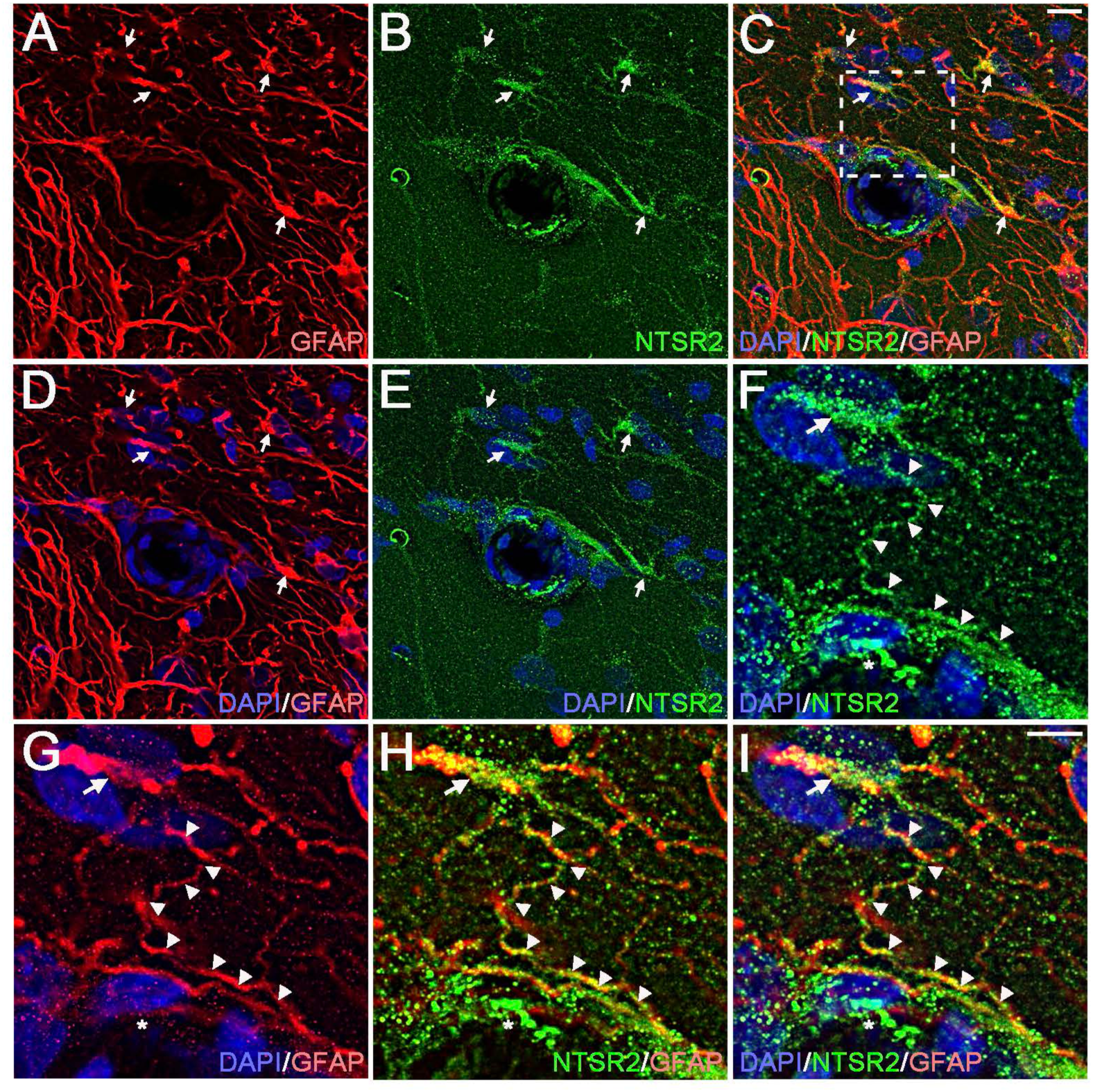
NTSR2 protein is expressed in astrocytic end feet around the blood vessels in the LM. GFAP (red, A,D,G) and NTSR2 (green, B,C,E,F,H,I) immunolabeling was performed in PILO 3D rats. All astrocytes (A-E, see arrows) around blood vessels expressed NTSR2. F-I show high magnification of the boxed-in area illustrating NTSR2 expression in the end-feet of astrocytes that project on blood vessels of the LM (arrowheads). In addition to astrocytes, other cells in and around the blood vessels expressed NTSR2. Cell nuclei were counterstained with DAPI (blue). Scale bars: 10 μm in all panels.

In order to determine the identity of the NTSR2-expressing cells in blood vessels, we used antibodies directed against endothelial and pericyte markers, namely the endothelial cell adhesion molecule, CD31, and the platelet derived growth factor receptor-beta, PDGFRβ, respectively (Uzunalli et al., 2019). The basal lamina was revealed using an antibody against Collagen IV (ColIV), which is the most abundant extracellular matrix protein in the endothelial basement membrane (Figs. 6-7).

**Figure 6.**
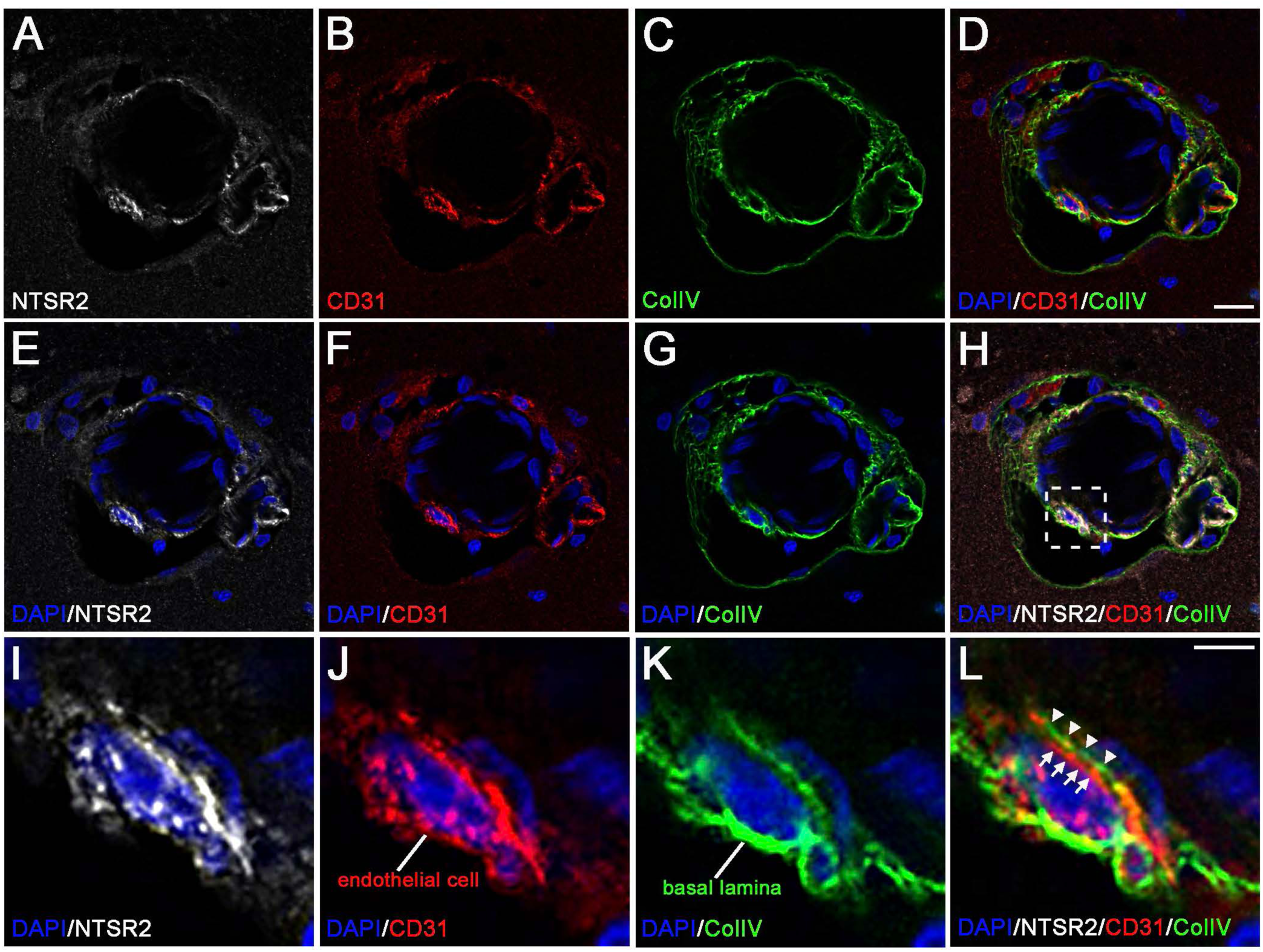
NTSR2 protein is expressed in blood vessel endothelial cells of the LM following pilocarpine-induced SE. NTSR2 (white), CD31 (red) and Collagen IV (green) immunostaining was performed in PILO 3D rats (A-H). I-L show high magnification of the boxed-in area illustrating endothelial cells (red, arrows) and basal lamina (green, arrowheads), revealed with anti-CD31 and -Collagen IV antibodies, respectively. NTSR2 labeling was mainly associated with endothelial cells as revealed with CD31 labeling. Cell nuclei were counterstained with DAPI (blue). Scale bars: 20 μm in A-H and 10 μm in I-L.

**Figure 7.**
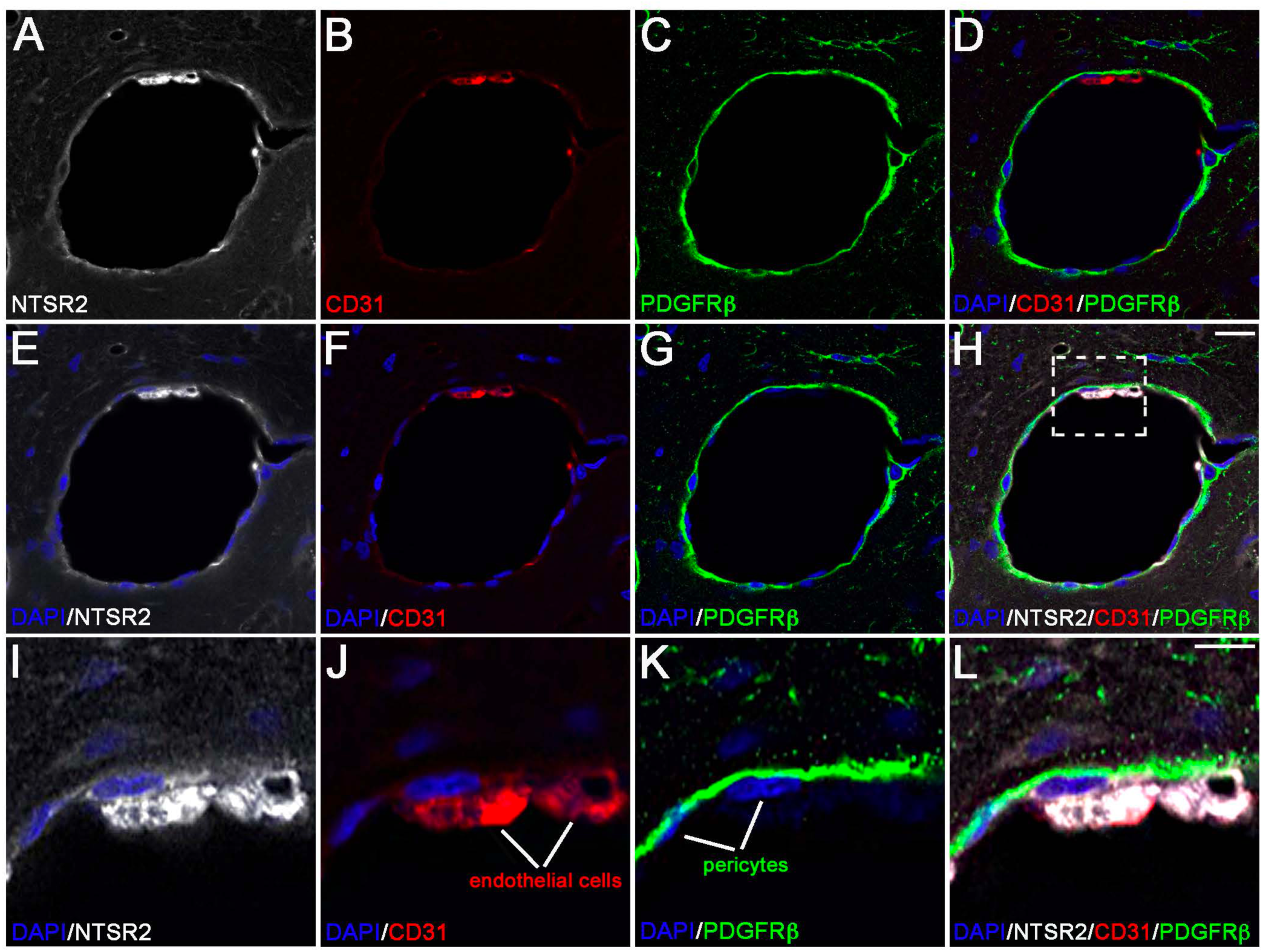
NTSR2 protein is virtually absent in blood vessel pericytes of the LM following pilocarpine-induced SE. NTSR2 (white), CD31 (red) and PDGFRβ (green) immunostaining was performed in PILO 3D rats (A-H). I-L show high magnification of the boxed-in area illustrating endothelial cells (red) and pericytes (green), revealed with anti-CD31 and -PDGFRβ antibodies, respectively. Co-labeling (NTSR2/CD31/PDGFRβ) confirmed NTSR2 expression in endothelial cells and faint or absent NTSR2 labeling in pericytes. Cell nuclei were counterstained with DAPI (blue). Scale bars: 20 μm in A-H and 10 μm in I-L.

We first examined whether endothelial cells expressed NTSR2. To this end, we performed NTSR2, CD31 and CollV triple immunolabeling in PILO 3D rats (Figs 6A-H). High magnification images of LM blood vessels revealed that NTSR2 was expressed in endothelial cells since both NTSR2 and CD31 stained the same cells (Figs. 6I,J). Endothelial cells displayed a punctiform NTSR2 immunolabeling pattern. Note that the ColIV immunolabeling (Fig. 6L, see arrowheads) was closely apposed to CD31 labeling (Fig. 6L, see arrows), as endothelial cells were surrounded by basal lamina. Finally, we sought to determine whether NTSR2 was expressed in blood vessel pericytes, which are present on the abluminal surface of the endothelium and in direct contact with endothelial cells while separated by the basement membrane (Armulik et al., 2011). NTSR2, CD31 and PDGFRβ triple immunolabeling was carried out in PILO 3D rats (Fig. 7). High magnification images showed that NTSR2 was faintly expressed in PDGFRβ-labeled pericytes, as compared with strong labeling of CD31-labeled endothelial cells (Figs 7I-L). The faint NTSR2 immunolabeling displayed by pericytes is more diffuse than that showed for astrocytes and endothelial cells. Thus, these results indicated that NTSR2 was expressed by several cellular components of the blood vessels.

Taken together, our data suggest that the increased NTSR2 expression in astrocytes and blood vessels in PILO rats is associated with neuroinflammation, since increased GFAP-labeled astrocytes were observed in the pilocarpine model. Finally, Iba1-labeled microglial cells around the blood vessels did not express NTSR2, similar to all other hippocampal areas (data not shown).

### Inflammation modulates differentially the expression of neurotensin receptors in glial primary cultures

Before investigating whether neuroinflammation can regulate NTSR2 expression, we first performed NTSR2 (Fig. 8B, green) and GFAP (Fig. 8C, red) double immunofluorescence labeling on primary glial cultures to validate our findings regarding NTSR2 astrocytic expression in rat brain tissue. Confocal microscopy analysis confirmed that all GFAP-labeled astrocytes were co-labeled for NTSR2 within the cytoplasm as well as at the cell membrane and processes (Figs. 8B,D, see arrowheads). The strong NTSR2 immunostaining observed in the cell body likely corresponds to the Golgi apparatus. We observed that not only astrocytes but also all cells in these glial cultures, that include astrocytes and microglial cells, were NTSR2-labeled. We performed NTSR2 (Fig. 8F, green) and Iba1 (Fig. 8G, red) double immunofluorescence on the same cultures and we report here for the first time that NTSR2 was expressed in microglial cells from primary glial cultures, in contrast to our *in vivo* findings. NTSR2 immunolabeling was more intense within the cell body and membrane compared to the cell processes (Figs. 8F,H). Surprisingly, we observed that expression of NTSR2 in microglia was stronger than that observed in astrocytes of CTL glial cultures. Indeed, quantitative analysis showed that the average fluorescence intensity for NTSR2 was significantly higher in Iba1-labeled microglia (149,4 ± 19,7%, 49%; *p*<0,05, Student’s *t*-test), compared with that of GFAP-labeled astrocytes (100 ± 13%).

**Figure 8.**
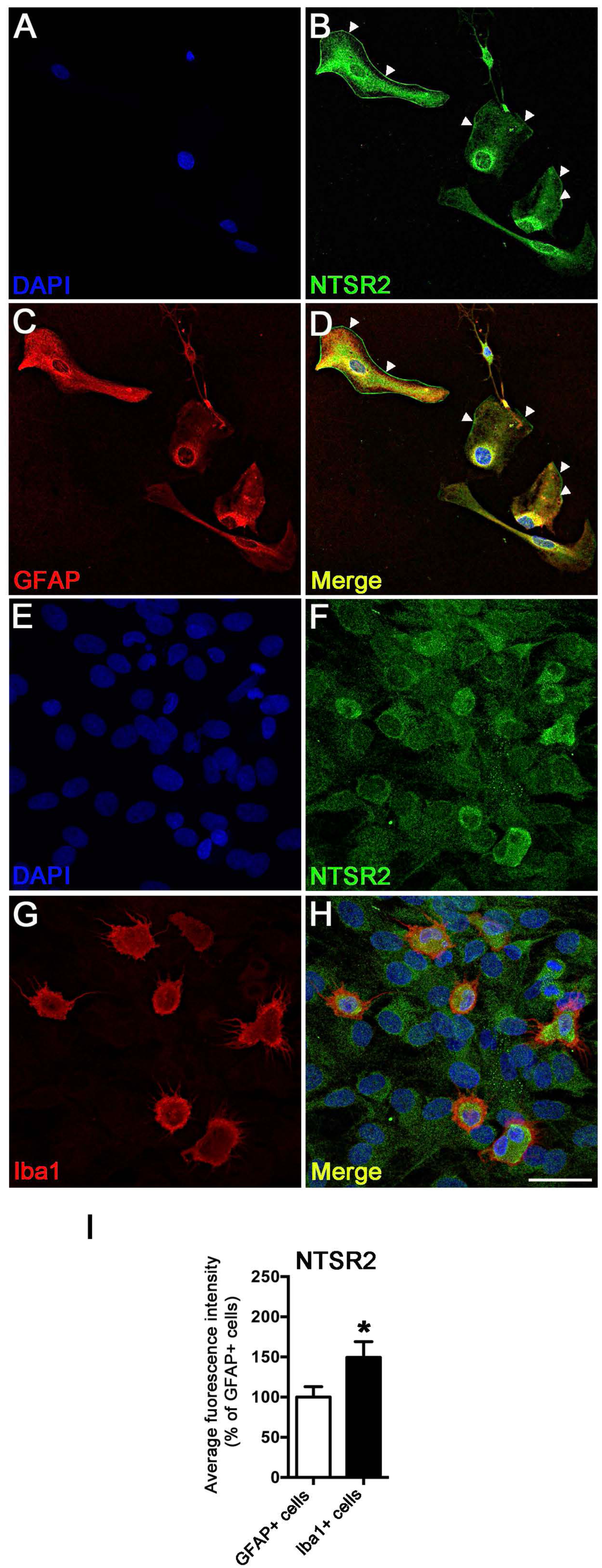
NTSR2 protein is expressed in primary cultured glial cells. A-D: Double NTSR2 (green, B) and GFAP (red, C) immunofluorescence shows that astrocytes from rat primary glial cultures express NTSR2 within the cell body and at the plasma membrane (see arrowheads). The strong immunostaining found in the cell body likely corresponds to the Golgi apparatus. E-H: NTSR2 (green, F) was also expressed in microglial cells from rat primary glial cultures, identified by Iba1 (red, G). Cell nuclei were counterstained with DAPI (blue, A,B,D & E,F,H). Merge corresponds to superimposition of DAPI, NTSR2 and GFAP (D), or DAPI, NTSR2 and Iba1 staining (H). I: The fluorescence intensity of NTSR2 labeling in CTL GFAP+ cells was measured and compared to that of CTL Iba1+ cells. The average intensity of NTSR2 was significantly higher in microglial cells compared to astrocytes (H). Values are given as the mean ± SEM as a percentage of the CTL GFAP+ cells. Asterisk indicates statistically significant difference: * *p<*0,05 (Student’s *t*-test). Scale bar: 10 μm

In order to study the effects of inflammation on the regulation of NTSR2 expression in primary glial cultures, cells were treated with different proinflammatory factors, including IL1β and LPS. The inflammatory response of these cells to IL1β or LPS was monitored by following the expression of IL1β and MCP-1/ccl2 mRNA by RT-qPCR at different time points (1, 6 and 24H), and synthesis of MCP-1/CCL2 by Elisa after 24H treatment. Quantitative analysis showed that IL1β did not affect the levels of IL1β mRNA at all time points analyzed (IL1β 1H: 1,4 ± 1,3; IL1β 6H: 3,2 ± 0,6; IL1β 24H: 3,3 ± 1,2) when compared to CTL levels (1 ± 0,0; n=8; *p*>0,05; Anova), whereas LPS significantly induced IL1β mRNA levels at 1H (8,6 ± 2,5) and 6H (11,5 ± 2,3). However, the levels of IL1β mRNA at 24H (3,6 ± 1,2) were not significantly different from CTL levels (1 ± 0,0; n=8; *p*>0,05; Anova) (Fig. 9A). The expression of MCP-1/ccl2 mRNA levels was significantly induced by IL1β as well as by LPS at 6H (IL1β 6H: 4,1 ± 1; n=8; *p*<0.05; LPS 6H: 8,2 ± 1; n=8; *p*<0,001, Tukey’s test) (Fig. 9B). However, no significant modulation of MCP-1/ccl2 mRNA was observed after either IL1β or LPS treatment at 1H (IL1β 1H: 1,6 ± 0,5; n=8; Anova; LPS 1H: 1,1 ± 0,2; n=8; *p*>0,05; Anova) and 24H (IL1β 24H: 2,4 ±1; n=8; *p*>0,05; Anova; LPS 24H: 2,8 ± 0,8; n=8; *p*>0,05; Anova) (Fig. 9B). Consistent with the RT-qPCR data, MCP-1/CCl2 is produced at significant levels in the supernatants of cultured glial cells after 24H treatment by IL1β (357,9 ± 25,4 ng/ml; 545,5%) or LPS (716 ± 27,4 ng/ml; 1091,3%) as compared to CTL (65,6 ± 14,1 ng/ml, 100%; n=8; *p*<0,01; Tukey’s test) (Fig. 9C). These data confirmed that our glial cultures were inflamed.

**Figure 9.**
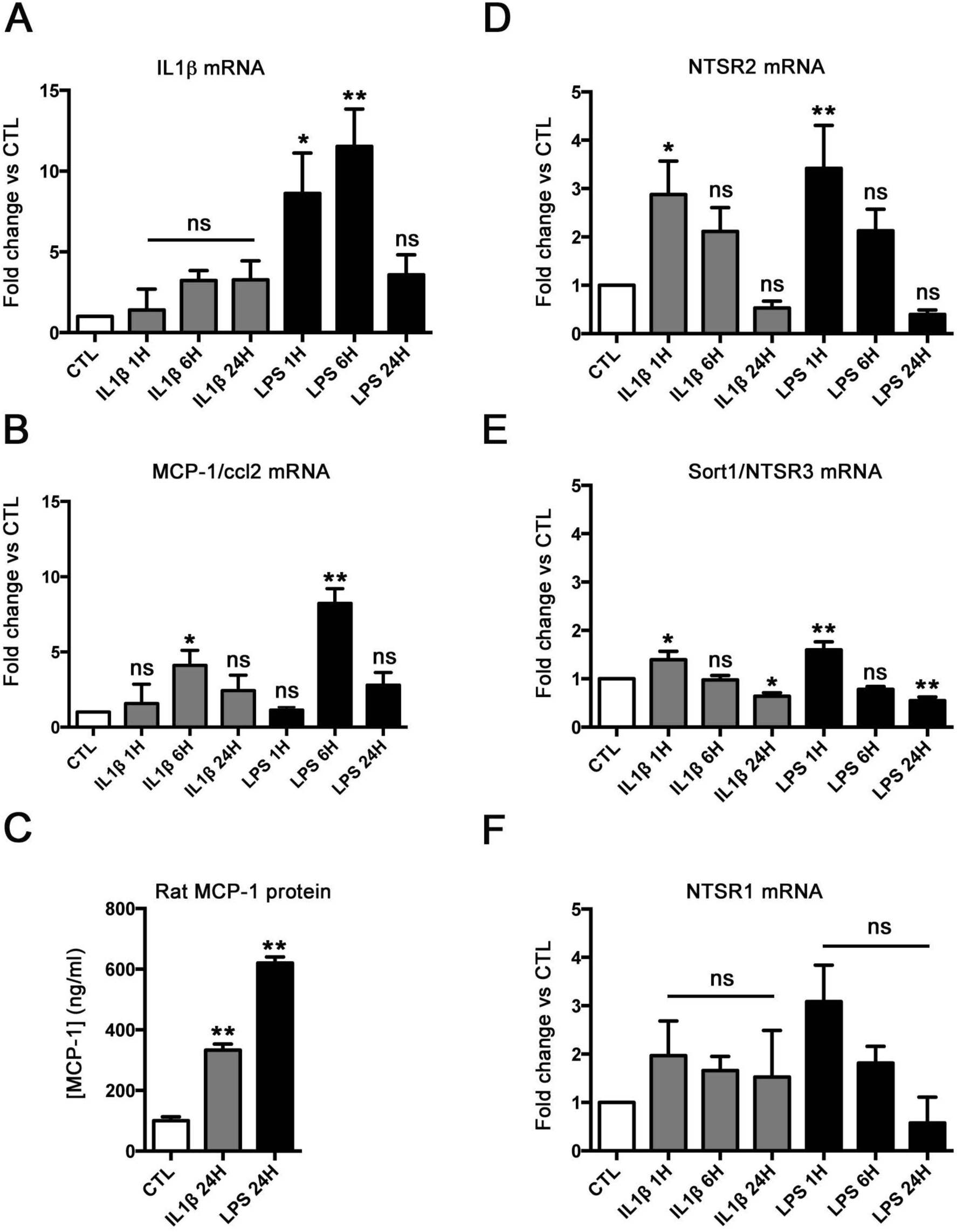
Inflammation differently affects the expression of neurotensin receptors in glial primary cultures. Histograms showing qRT-PCR quantification of the inflammatory cytokines IL1β (A) and MCP-1/ccl2 mRNA levels (B), after 1, 6 and 24H treatment with either IL1β (10 ng/ml) or LPS (1 μg/ml), and MCP-1/CCL2 protein in the supernatants of cultured glial cells assessed by ELISA (C), after 24H treatment with either IL1β or LPS. IL1β mRNA was induced after 1 and 6H treatment with LPS, and MCP-1/ccl2 mRNA was induced after 6H treatment with either IL1β or LPS. As a result, MCP-1/CCL2 protein was increased in primary glial cultures after IL1β or LPS treatment. D, E, F: Histograms showing qRT-PCR levels of mRNAs encoding NTSR2 (D), and the other NT receptors, Sortilin1/NTSR3 (Sort1, E) and NTSR1 (F), after 1, 6 and 24H treatment with either IL1β or LPS. NTSR2 and NTSR3 mRNAs were induced after 1H treatment with IL1β or LPS, while NTSR1 mRNA levels were not modulated. These results show that NTSR2 and NTSR3 are upregulated in primary glial cells during inflammation and that inflammation modulated differentially the expression levels of the NT receptors. Values for A, B, D, E, F are given as the mean ± SEM normalized to CTL. Asterisks indicate statistically significant differences: * *p<*0,05, ** *p<*0,01 (one-way ANOVA followed by Tukey’s *post hoc* test).

We next analyzed the expression of NTSR2 and the other receptors, NTSR1 and NTSR3, after 1, 6 and 24H treatment with either IL1β or LPS. Inflammation induced rapid upregulation of mRNAs encoding NTSR2 (IL1β 1H: 2,9 ± 0,7; n=8; *p*<0,05; Tukey’s test; LPS 1H : 3,4 ± 0,9; n=8; *p*<0,01; Tukey’s test) (Fig 9D) and to a lower extent NTSR3 (IL1β 1H: 1,4 ± 0,2; n=8; *p*<0,05; Tukey’s test; LPS 1H : 1,60 ± 0,2; n=8; *p*<0,001; Tukey’s test) (Figs 9D,E) at 1H after IL1β or LPS treatments, whereas no significant modulation of NTSR2 (IL1β 6H: 1,4 ± 0,5; n=8; *p*>0,05; Anova; LPS 6H: 2,1 ± 0,4; n=8; *p*>0,05; Anova) and NTSR3 (IL1β 6H: 1 ± 0,1; *p*>0,05; Anova; LPS 6H: 0,8 ± 0,1; n=8; *p*>0,05; Anova) mRNA was observed after either IL1β or LPS treatment at 6H (Figs. 9D,E). However, significant decrease of mRNA encoding NTSR3 was observed at 24H after IL1β or LPS treatments (IL1β 24H: 0,6 ± 0,1; n=8; *p*<0,05; Tukey’s test; LPS 24H: 0,5 ± 0,1; n=8; *p*<0,01; Tukey’s test) (Fig 9E). Our study showed no significant modulation of NTSR1 encoding mRNA at all time points examined (IL1β 1H: 2 ± 0,7; IL1β 6H: 1,7 ± 0,3; IL1β 24H: 1,5 ± 0,8; LPS 1H: 3,1 ± 0,8; LPS 6H: 1,8 ± 0,4; LPS 24H : 0,6 ± 0,2; n=8; *p*>0,05; Anova) (Fig. 9F). These results indicate that inflammation differently affects the expression of neurotensin receptors in primary glial cultures.

### NTSR2 is expressed as an immediate early gene following inflammatory stimuli in glial primary cultures

The transient and rapid (as early as 1H) upregulation by inflammation of mRNAs encoding NTSR2 and NTSR3 was reminiscent of the pattern of expression of immediate-early genes (IEGs), also referred to as primary response genes (Herschman, 1991; Lau & Nathans, 1991). IEGs are also characterized by the fact that the transient and rapid response to stimuIi does not require protein synthesis, and that their expression is usually induced by agents that block protein synthesis (Mehendale & Apte, 2009). In order to determine whether the activation of NTSR2 and NTSR3 requires protein synthesis, we incubated glial cells with or without LPS (1 μg/ml) and/or cycloheximide (CHX), a protein synthesis inhibitor (Alberini, 2008), at a classical dose of 10 μg/ml (Cochran et al., 1984; Greenfield et al., 1996), for 1, 6 and 24H prior to RNA extraction and RT-qPCR analysis. We used as a standard positive control Zif-268/egr-1, a zinc-finger protein known to be rapidly induced in diverse cell types following activation by a variety of stimulants, including mitogens, cell differentiating agents, and LPS (Gashler & Sukhatme, 1995; Yao et al., 1997; Xu et al., 2001). We first evaluated whether Zif-268/egr-1 behaved as an IEG in primary rat glial cells. RT-qPCR analysis demonstrates that in these cells, LPS alone induced strong, rapid and transient expression of Zif-268 mRNA, with a peak in mRNA levels occurring at the 1H time point (23 ± 0,5; 23-fold; n=3; *p*<0,001; Tukey’s test) as compared to untreated CTL cells (1 ± 0,0) (Fig.10A). Zif-268 mRNA expression levels returned back to baseline at the 6 and 24H time points (Fig. 10 A, left plot), consistent with the temporal profile of *zif-268* gene induction described previously (Hughes et al., 1993; Yamaji et al., 1994). CHX alone resulted in a more potent induction of Zif-268/egr-1 mRNA at all time points compared to CTL and LPS (Fig. 10, right plot) with a 60- fold induction at 1H (59,9 ± 2; n=3; *p*<0,01; Tukey’s test) compared to CTL (Fig. 10A, right diagram) while CHX alone resulted in a 2,6-fold induction at 1H (59,9 ± 2; n=3; *p*<0,01; Tukey’s test) when compared to LPS 1H (23 ± 0,5). When combining LPS with CHX in glial cell cultures, we observed a super-induction of Zif-268/egr-1 mRNA at all time points compared to LPS (Fig. 10A, left plot) with a 7,7-fold super-induction of Zif-268/egr-1 mRNA levels at 1H (177,1 ± 15,3; n=3; *p*<0,05; Tukey’s t-test) over LPS alone (Fig. 10A, right diagram). This LPS-CHX super-induction was most obvious when the Zif-268/egr-1 mRNA levels returned back to CTL levels following LPS treatment (Fig. 10A, left plot). These data suggest that in rat primary glial cells, inhibition of translation prevents the synthesis of a protein/or proteins that negatively regulate Zif-268/erg-1 expression. Thus, the *zif-268* gene is induced by LPS in rat primary glial cells and this induction is independent of *de novo* protein synthesis. These results show that Zif-268 is induced as an IEG in primary glial cultures activated by LPS, consistent with other cell types (Hughes et al., 1993; Yamaji et al., 1994; Gashler et al., 1995; Yao et al., 1997; Xu et al., 2001).

**Figure 10.**
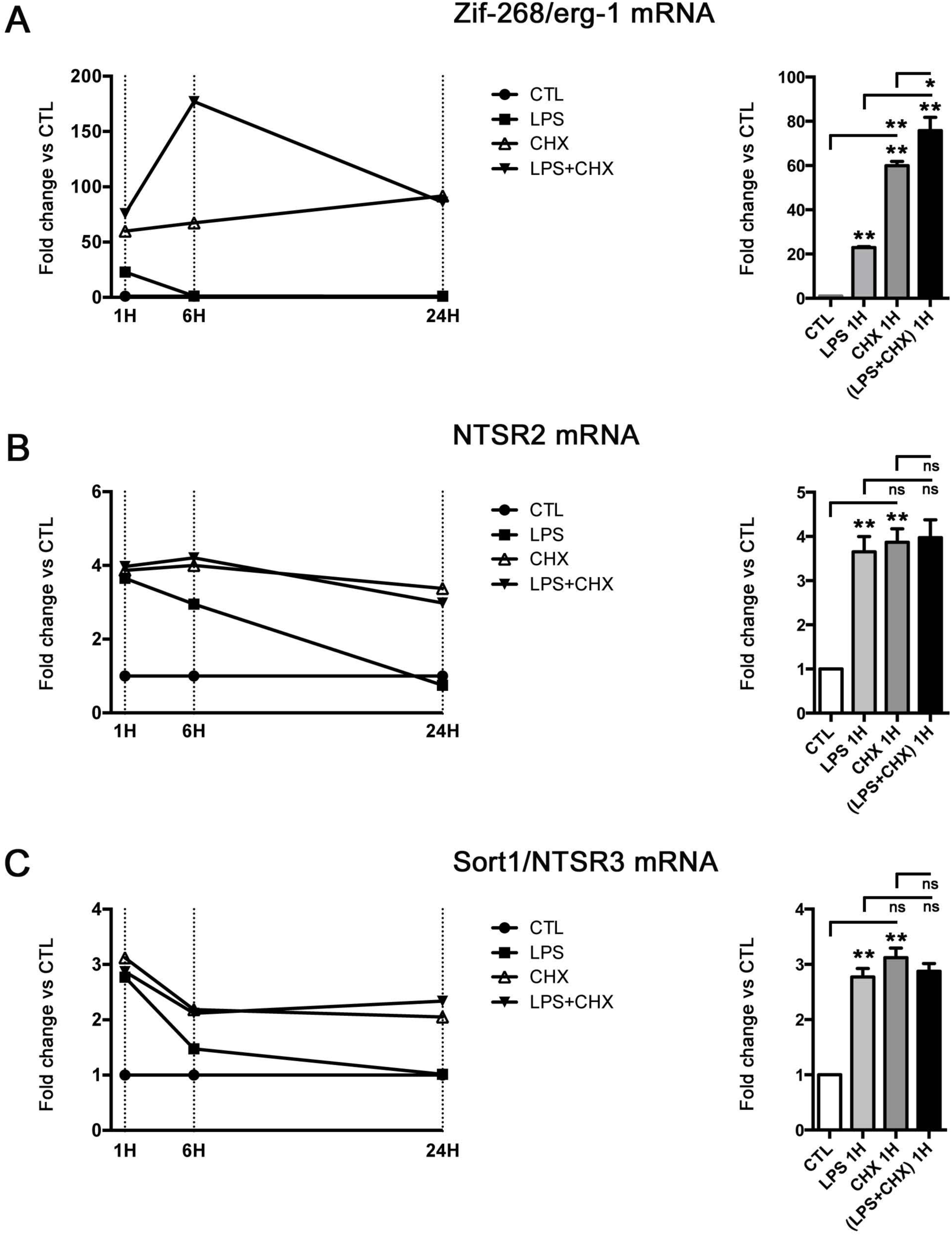
The rapid induction by inflammation of mRNAs encoding neurotensin receptors is independent of *de novo* protein synthesis in glial primary cultures. Plots on the left side illustrate the modulation of mRNA levels of the Zif-268/erg-1 IEG (A) and neurotensin receptors, NTSR2 (B) and Sort1/NTSR3 (C) at 1, 6 and 24H after treatment with LPS, CHX or LPS + CHX. LPS alone induced moderate to strong, rapid and transient expression of Zif-268/egr-1, NTSR2 and Sort1/NTSR3 mRNA with a peak occurring at the 1H time point when compared to untreated glial cells. Histograms on the right side illustrate Zif-268/erg-1, NTSR2 and Sort1/NTSR3 mRNA expression after 1H treatment with LPS, CHX or LPS + CHX. Treatment with cycloheximide (CHX, 10 μg/ml) induced the expression of Zif-268/erg-1, compared to untreated CTL glial cells and LPS treatment. However, CHX alone did not affect the expression of NTSR2 and Sort1/NTSR3 compared to LPS. When LPS was combined to CHX, super-induction was observed for Zif-268/erg-1 but not for NTSR2 and Sort1/NTSR3 as compared to LPS and CHX. Thus, as for IEG Zif-268/egr-1, the rapid and transient kinetics of activation and the absence of inhibition of NTSR2 and NTSR3 mRNA expression by CHX, suggest that NTSR2 and NTSR3 are new IEGs in rat primary glia. Values are given as the mean ± SEM normalized to CTL. Asterisks indicate statistically significant differences: * *p<*0,05, ** *p<*0,01 (one-way ANOVA followed by Tukey’s *post hoc* test).

As shown in Figure 9D,E, LPS alone also induced rapid and transient expression of NTSR2 and NTSR3 (Figs. 10B,C, left plots), with mRNA expression peaking at the 1H time point (NTSR2: 3,7 ± 0,3; NTSR3: 2,8 ± 0,2; n=3; *p*<0,001; Tukey’s test) when compared to untreated CTL glial cells (1 ± 0,0) (Figs. 10B,C, right diagrams). LPS induction of NTSR2 (3,7-fold) and NTSR3 (2,8-fold) mRNA levels was less potent than that of Zif-268 (23-fold), and decreased less rapidly (Figs. 10B,C, left plots). CHX-treatment alone also induced NTSR2 (3,9 ± 0,3; 6H: 3 ± 0,2; 3-fold; 24H: 3,4 ± 0,2; 3,4-fold; n=3; *p*<0,001; Tukey’s t-test) and NTSR3 (1H: 3,1 ± 0,2; 3,1-fold; 6H: 2,2 ± 0,1; 2,2-fold; 24H: 2,1 ± 0,0; 2,1-fold; n=3; *p*<0,001; Tukey’s test) mRNA levels at all time points compared to CTL (Figs. 10A-C), but to a lesser extent than the Zif-268/erg-1 induction by CHX (from 60-fold induction at 1H to 91,8-fold at 24H) (Fig. 10 A). In contrast to Zif-268/erg-1, NTSR2 and NTSR3 did not show any significant super-induction response at all time points examined when CHX treatment was combined with LPS or CHX treatment alone (Figs. 10B,C, left plots and right diagrams). Thus, although CHX induced expression of NTSR2 and NTSR3 in glial cells, there was no additional stimulation of these genes by LPS treatment (Figs. 10B,C, left plots). CHX treatment did not block activation of NTSR2 and NTSR3, similarly to what was described for c-fos (Yamaji et al., 1994) and BDNF (Hughes et al., 1993). Our results show that upon stimulation with a proinflammatory factor such as LPS, induction of NTSR2 and NTSR3 mRNA levels were not dependent upon *de novo* protein synthesis. Taken together with the rapid and transient kinetics of activation, they suggest that NTSR2 and NTSR3 are new IEGs in rat glial cells.

### The vectorized NT analog VH-N412 regulates NTSR2, GFAP and Iba1 protein expression in primary glial cultures

We developed VH-N412 is a vectorized version of a NT functional fragment. It encompasses the C-terminal NT part (8-13, RRPYIL-OH), the shortest NT fragment with full binding and pharmacological activities (Granier et al., 1982) conjugated via a PEG6 linker to VH4129, a cyclic peptide ([cMPipRLRSarC]c) encompassing non-natural amino-acids, that binds to the low-density lipoprotein receptor (LDLR) and that we previously described (Jacquot et al., 2016). VH4129 belongs to a family of cyclic peptides that can be conjugated to a variety of cargos including fluorophores, peptides and proteins to promote transcytosis across the BBB and enhance their cellular uptake and brain delivery that we previously described (Malcor et al., 2012; Jacquot et al., 2016; Molino et al., 2017; David et al., 2018), and that others have used to functionalize nanoparticles and liposomes to increase brain delivery of anticancer drugs (Zhang et al., 2013; Chen et al., 2017; Shen et al., 2018). VH-N412 undergoes transcytosis across the BBB and induces rapid hypothermia when administered i.v. to rats or mice, and we have shown that it elicits neuroprotective and anti-inflammatory effects following *in vivo* administration in KA-induced TLE in mice and neuroprotection *in vitro* (Soussi et al., manuscript in preparation). Since NTSR2 mRNA expression is regulated by inflammation, we investigated whether its vectorized ligand VH-N412 could in turn modulate NTSR2 protein expression in primary inflamed glial cultures. To this end, dual NTSR2 (green) and GFAP (red) immunocytochemistry (Fig. 11) was carried out on primary glial cultures treated with 1 μM VH-N412 (Figs. 11b,h,n), 10 ng/ml IL1β (Figs. 11c,i,o), 10 ng/ml IL1β + 1 μM VH-N412 (Figs. 11d,j,p), 10 μg/ml LPS (Figs. 11e,k,q) or 10 μg/ml LPS + 1 μM VH-N412 (Figs. 11f,l,r) for 24 H, and compared to untreated glial cells (Figs 11a,g,m). The dose of 1 μM was chosen based on neuroprotection effects obtained at this dose compared to 0,1, 1, and 10 μM (Soussi et al., manuscript in preparation). Quantitative analysis showed that 1 μM VH-N412 alone did not modulate NTSR2 protein expression (93,7 ± 13,7; n=3; p>0,05; Anova), as compared to untreated CTL astrocytes (100 ± 13) (Fig. 11B). IL1β caused significant increase of NTSR2 protein expression (199,1 ± 38,5; 99%; n=3; p<0,05; Tukey’s test) compared to CTL, which was significantly down-regulated upon VH-N412 treatment (IL1β + VH-N412: 66,4 ± 9,3; 133%; n=3; p<0,01; Tukey’s test). LPS alone or together with VH-N412 did not modulate NTSR2 protein expression (LPS: 150 ± 29,6; LPS + VH-N412: 91,8 ± 14,7; n=3; p>0,05; Anova) (Fig. 11B). In contrast to NTSR2, VH-N412 alone significantly reduced GFAP protein expression (51 ± 3,6; n=3; p<0,05; Tukey’s test) compared to untreated CTL cultures (100 ± 13) (Fig. 11C), suggesting that VH-N412 reduced astrocytic inflammation. IL1β alone (102 ± 10; n=3; p>0,05; Anova) and IL1β + VH-N412 (110,9 ± 16,4; n=3; p>0,05; Anova) had no significant effect on GFAP protein levels compared to untreated CTL astrocytes (Fig. 11C). GFAP levels were not modulated by LPS (132,8 ± 13,8; n=3; p>0,05; Anova) when compared to CTL untreated astrocytes (Fig 11C). However, VH-N412 significantly reduced GFAP expression in the LPS-treated astrocytes (LPS + VH-N412: 75,7 ± 6,1; 57%; n=3; p<0,01; Tukey’s test) (Fig 11C). Altogether, our data indicate that VH-412 may regulate astrocytic inflammation.

**Figure 11.**
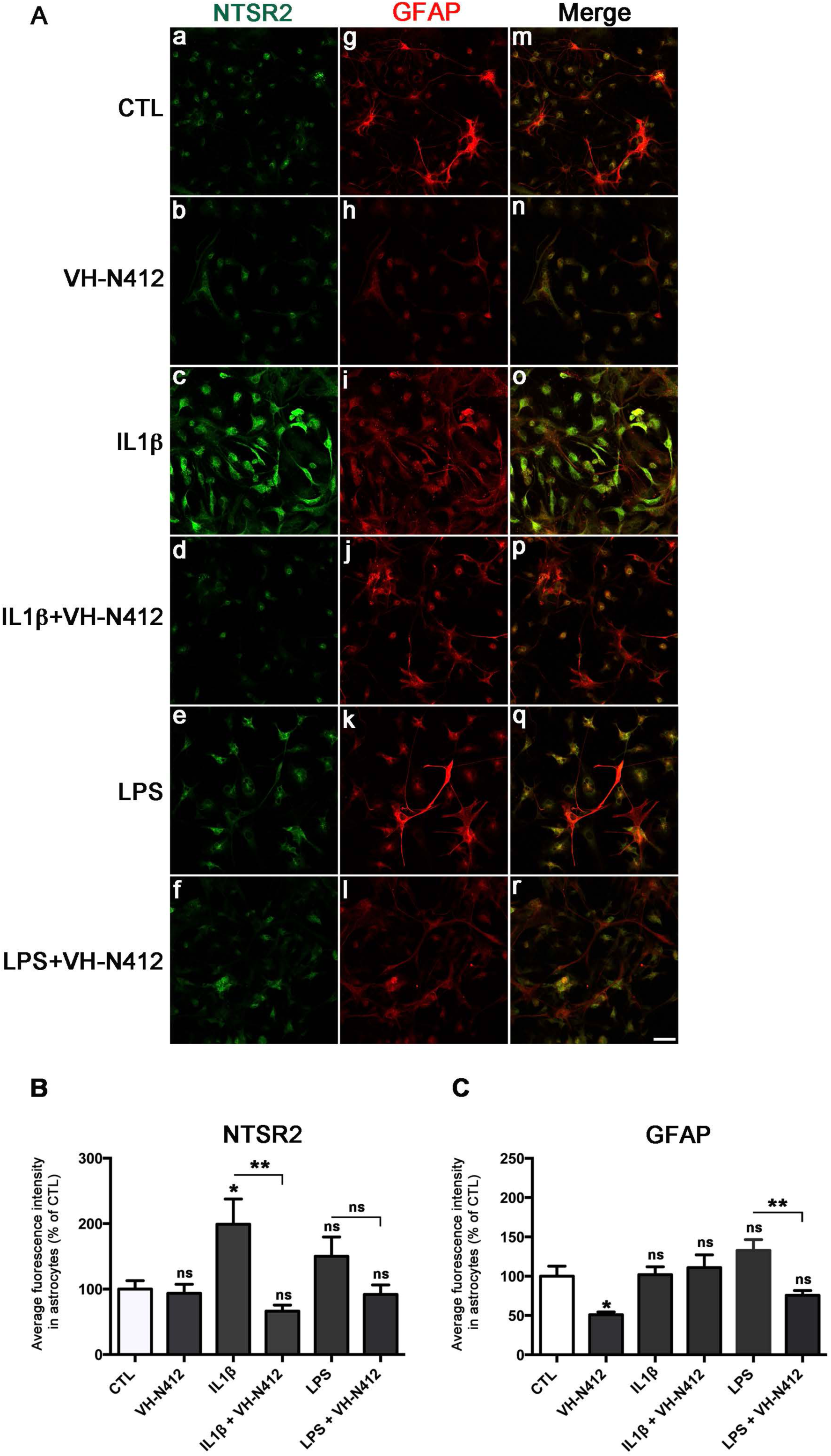
The vectorized NT analog VH-N412 down-regulates NTSR2 and GFAP protein expression in primary glial cultures. Representative images of NTSR2 and GFAP staining in untreated primary cultures of glial cells (CTL) or after 24H treatment with VH-N412 (1 μM), IL1β, IL1β + VH-N412, LPS, or LPS + VH-N412 (A). Merge corresponds to superimposition of NTSR2 and GFAP staining. In (B), histograms comparing the average fluorescence intensity of NTSR2 in astrocytes after treatment with VH-N412, IL1β, IL1β + VH-N412, LPS, or LPS + VH-N412 compared to untreated CTL astrocytes, and comparison between the different treatments. Quantitative analysis showed that the NT analogue VH-N412 significantly decreased NTSR2 protein expression to CTL levels after 24H IL1β treatment in GFAP+ astrocytes. However, VH-N412 did not affect NTSR2 protein expression after 24H LPS treatment in GFAP+ astrocytes. In (C), histograms comparing the average fluorescence intensity of GFAP in astrocytes after treatment with VH-N412, IL1β, IL1β + VH-N412, LPS, or LPS + VH-N412, compared to untreated CTL astrocytes, and comparison between the different treatments. Quantitative analysis showed that VH-N412 alone significantly decreased GFAP protein expression compared to untreated CTL astrocytes. However, VH-N412 did not affect GFAP protein expression after 24H IL1β treatment in GFAP+ astrocytes. Finally, VH-N412 significantly decreased GFAP protein expression back to CTL levels after 24H LPS treatment in GFAP+ astrocytes compared to LPS alone-treated cells. Values are given as the mean ± SEM as a percentage of CTL. Asterisks indicate statistically significant differences: * *p<*0,05, ** *p<*0,01 (one-way ANOVA followed by Tukey’s *post hoc* test).

Considering that NTSR2 is expressed also in microglial cells *in vitro*, NTSR2 and Iba1 immunocytochemistry (Fig. 12) was carried out on the same glial cultures treated under the same conditions: 1 μM VH-N412 (Figs. 12b,h,n), 10 ng/ml IL1β (Figs. 11c,i,o), 10 ng/ml IL1β + 1 μM VH-N412 (Figs. 12d, j, p), 10 μg/ml LPS (Figs. 12e, k, q) or 10 μg/ml LPS + 1 μM VH-N412 (Figs. 12f, l, r) for 24 H. Treated microglia were compared to untreated microglia (Figs 12a,g,m) and the levels of NTSR2 and Iba1 were quantified. As for astrocytes, VH-N412 alone did not regulate NTSR2 protein expression (102,1 ± 7,9; n=3; *p*>0,05; Anova) in microglial cells, as compared to CTL untreated microglia (100 ± 13,2) (Fig. 12B). IL1β tended to increase NTSR2 protein levels but did not reach the level of significance (105,5 ± 12; n=3; *p*>0,05; Anova), as compared to those of CTL. On the other hand, LPS led to significant increased NTSR2 protein expression (189,7 ± 26,2; 90%; n=3; *p*<0,01; Tukey’s test). Treatment of the cultures with VH-N412 simultaneously with IL1β or LPS led to significantly decreased NTSR2 levels compared to treatment, either with IL1β or LPS alone (IL1β + VH-N412: 47,1 ± 7,8; 104%; LPS + VH-N412: 63,3 ± 4,4; 127%; n=3; *p*<0,01; Tukey’s test) (Fig. 12B). VH-N412 treatment in our non-inflamed primary glial cultures did not result in changes of Iba1 protein expression (78,1 ± 6,8; n=3; *p*>0,05; Anova), as compared to CTL (100 ± 9,6) (Fig. 12C). However, Iba1 protein expression was significantly increased after 24H treatment with either IL1β (183,7 ± 16; 84%; n=3; *p*<0,05; Tukey’s test) or LPS (266,7 ± 35,2; 167%; n=3; *p*<0,01; Tukey’s test). Finally, VH-N412 significantly decreased Iba1 protein expression both in the IL1β (53,9 ± 6,6; 130%; n=3; *p*<0,01; Tukey’s test) and in the LPS-treated microglia (109,9 ± 8,6; 157%; n=3; *p*<0,01; Tukey’s test), suggesting its potential role in modulating microglial inflammation. Collectively, these results show that NTSR2 is induced at high levels in reactive astrocytes and microglia during pathology, and that astrocyte and microglial reactivity can be modulated by NT analogues.

**Figure 12.**
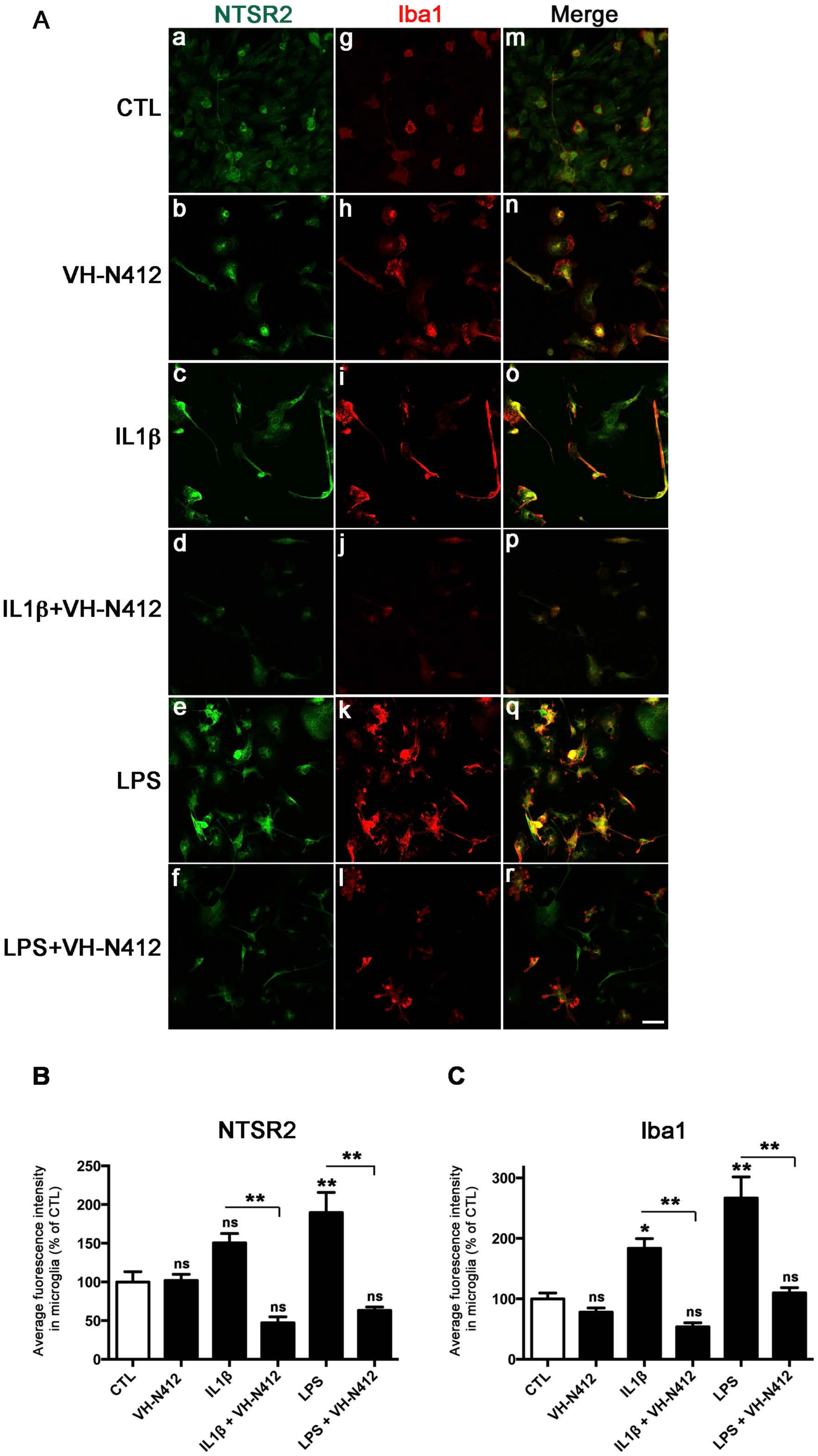
The vectorized NT analog VH-N412 down-regulates NTSR2 and Iba1 protein expression in primary glial cells. Representative images of NTSR2 and Iba1 staining in untreated primary cultures of glial cells (CTL) or after 24H treatment with VH-N412 (1 μM), IL1β, IL1β + VH-N412, LPS, or LPS + VH-N412 (A). Merge corresponds to superimposition of NTSR2 and Iba1 staining. In (B), histograms comparing the average fluorescence intensity of NTSR2 in Iba1-positive microglia after treatment with VH-N412, IL1β, IL1β + VH-N412, LPS, or LPS + VH-N412, compared to untreated CTL Iba1-positive microglia, and comparison between the different treatments. Quantitative analysis showed that VH-N412 significantly decreased NTSR2 protein expression back to CTL levels after 24H IL1β or LPS treatment in Iba1+ microglia. In (C), histograms comparing the average fluorescence intensity of Iba1 in microglia after treatment with VH-N412, IL1β, IL1β + VH-N412, LPS, or LPS + VH-N412, compared to untreated CTL astrocytes, and comparison between the different treatments. Quantitative analysis showed that VH-N412 significantly decreased Iba1 protein expression back to CTL levels after 24H IL1β or LPS treatment in Iba1+ microglia. Values are given as the mean ± SEM as a percentage of CTL. Asterisks indicate statistically significant differences: * *p<*0,05, ** *p<*0,01 (one-way ANOVA followed by Tukey’s *post hoc* test).

## DISCUSSION

Our study shows that NTSR2 protein is expressed in astrocytes of the adult rodent hippocampus and its expression is increased together with astrocyte reactivity in the hippocampus at the early phases following induction of SE, from 3 to 14 days, and decreased at 3 months post-SE. All the GFAP positive cells co-expressed NTSR2. Following SE, NTSR2 immunoreactivity was also increased in perivascular astrocytes and their end-feet and was apparent in endothelial cells. Proinflammatory factors such as IL-1β and LPS induced NTSR2 in astrocytes, but also in microglia *in vitro*. Astrocytic NTSR2 expression showed characteristic immediate early gene response under inflammatory conditions. Treating cultured glial cells with a vectorized NT analogue decreased NTSR2, GFAP and Iba1 expression, suggesting a role for NTSR2 in the downregulation of astroglial neuroinflammation by NT.

### NTSR2 protein is expressed in astrocytes in rat brain

In the present work, we investigated *in vivo* and *in vitro*, by immuno-histo- and immuno-cytochemistry laser scanning confocal microscopy approaches, the expression of the NTSR2 protein in rat hippocampal cells using GFAP labeling combined with a specific and validated anti-NTSR2 antibody. Our study clearly demonstrates for the first time that the NTSR2 protein is expressed in the DG and CA1-CA3 regions of Ammon’s horn.

Recently, NTSR2 was shown to be expressed in the DG and CA3 in a transgenic mice line (NtsR2^Cre/+,GFP^), however the identity of the non-neuronal (NeuN-negative) NTSR2-expressing cells was not revealed (Manning et al., 2019). In our study, we observed differential NTSR2 expression in hippocampal neurons (Kyriatzis et al., in preparation) but in the present report we chose to focus our attention on the non-neuronal expression of the NTSR2 protein, in particular in astrocytes. We show for the first time NTSR2 protein expression in all GFAP-labeled astrocytes. A high throughput study showed that NTSR2 mRNA is one of the most abundant receptor-encoding mRNAs in purified mice astrocytes (Y. Zhang et al., 2014), and *in vitro,* NTSR2 has been used as an astrocytic marker in single-cell transcriptomic analysis (Dulken et al., 2017). RT-qPCR assays showed that cultured astroglial cells express NTSR2 mRNA and its vNTSR2 isoform rather than NTSR1 (Ayala-Sarmiento et al., 2015). *In situ* hybridization studies *in vivo* showed NTSR2 mRNA expression in rat astrocytes (Walker et al., 1998; Yamauchi et al., 2007), and in mouse astrocytes of P17 mice (Cahoy et al., 2008). Dual immunohistochemical labeling of GFAP and *in situ* hybridization of NTSR2 mRNA suggested that NTSR2 was expressed only a small subset of astrocytes in the rat brain (Nouel et al., 1999). More recently, *in situ* hybridization localized NTSR2 mRNA in a subpopulation of astrocytes of the median preoptic nucleus (Tabarean, 2020). At the protein level, previous studies based on NT-binding experiments and sensitivity to levocabastine, a selective non-peptide histamine H1 receptor antagonist and NTSR2 ligand that does not bind to NTSR1 nor NTSR3 (Sarret & Beaudet, 2002), suggested NTSR2 protein expression in cultured astrocytes; bound fluorescent NT indicated that receptor expression concerned only a sub-population of astrocytes in culture. More recently, NTSR2 immunostaining was reported in astrocytes of the ventral tegmental area (Woodworth et al., 2018). However, other studies based on double-immunolabeling combining an N-terminal-specific anti-NTSR2 antibody and the astrocyte marker calcium-binding protein S100β, reported absence of NTSR2 immunostaining in adult rat brain astrocytes (Sarret et al., 2003).

With regard to subcellular localization, we observed *in vivo* but even more clearly *in vitro*, that NTSR2 is localized at the cell membrane, typical for a receptor; however we also observed NTSR2 aggregates in astrocytic processes and cell bodies, reminiscent of NTSR2 localization in intracellular vesicles and in the *trans*-Golgi network described in rat spinal cord neurons (Perron et al., 2006).

### NTSR2 is induced in reactive astrocytes in rodent models of epilepsy

We questioned whether astrocytic NTSR2 expression was modulated in pathological situations involving astrocyte reactivity. To address this issue, we studied two pathophysiological models of TLE induced by pilocarpine in rats and KA in mice. NTSR2 expression was largely increased both after pilocarpine- and KA-induced SE in all hippocampal areas. The NTSR2 increase following SE coincides with the peak of astrocyte activation and prolonged elevation of astrocytic Ca^2+^ signaling, which occurs at 3 days post-SE in different rodent experimental models (Seifert et al., 2010). These reactive astrocytes were shown to contribute to delayed neuronal death of cortical and hippocampal pyramidal cells following pilocarpine-induced SE (Ding et al., 2007). Accordingly, at 3 months post-SE, characterized by spontaneous recurrent seizures and attenuated astrocytic reactivity (Garzillo and Mello, 2002; Choi and Koh, 2008), we observed concomitant decrease of NTSR2. The fact that the NTSR2 expression pattern closely follows that of astrocyte reactivity during the course of the disease, provides a hint that this receptor is involved in the neuroinflammation processes associated with pathology. Despite well-established commonalities, astrocyte reactive gliosis can be heterogeneous in response to specific injury. For instance, the molecular phenotypes of reactive astrocytes induced by ischemia and LPS suggest that they may be beneficial or detrimental, respectively (Zamanian et al., 2012). In other neuroinflammatory settings, such as those induced by brain stab-wound in rats, a marked increase in the number of NTSR2 mRNA expressing astrocytes around the lesion has been described, associated with increased NTSR2 mRNA expression at the cellular level (Nouel et al., 1999). Taken with our data, these results suggest that NTSR2 upregulation during astrocytic reaction may be common to different brain injuries. Although we did not observe NTSR2 labeling in CTL mouse MAP2-negative cells resembling astrocytes, our immunohistochemical analysis of mice brain after KA injection also showed enhanced expression of NTSR2 protein following acute neuronal injury. The fibrillose appearance of the labeled cells and absence of co-labeling with the mature neuronal marker MAP2 suggest they are astrocytes. Thus, NTSR2 upregulation in astrocytes seems to be common to different species and to different pathophysiological conditions inducing TLE.

### NTSR2 is induced in reactive astrocytes and microglial cells *in vitro* following proinflammatory stimulation

A major finding of our study is that inflammation mediated by proinflammatory factors IL1β and LPS significantly enhanced both NTSR2 mRNA and protein levels in astroglial cells, congruent to our *in vivo* data. This is the first demonstration of NTSR2 mRNA and protein up-regulation following IL1β-mediated inflammation and results are in agreement with Pang et al. (2001), who reported increased expression of NTSR2 mRNA in purified astrocytes following LPS stimulation. In the adult rat hippocampus, we showed that NTSR2 was expressed specifically in astrocytes and not in CTL or reactive microglia, that also contribute to inflammation associated with epilepsy. In the pilocarpine model, we observed strong Iba1 immunolabeling, indicative of microglial reactivity. However, we found no NTSR2 staining in Iba1-positive cells *in vivo*, despite the fact that we observed strong NTSR2 expression in microglial cells *in vitro*. This discrepancy could result from insufficient microglia activation in the pilocarpine model of epilepsy, raising the question whether in another pathological model, appropriate proinflammatory factors could induce NTSR2 expression in activated microglia. Alternative hypotheses include differential behavior of microglial cells in *in vivo* and *in vitro* conditions, and/or distinct microglia responses to the proinflammatory factors expressed *in vivo* and those we assessed *in vitro*. To our knowledge, this is the first demonstration of NTSR2 expression in microglia, as previous studies reported that murine and human microglia cells express only NTSR3 but not NTSR1 nor NTSR2 (Martin et al., 2003; Patel et al., 2016).

### VH-N412 decreases astrocyte and microglial reactivity

*In vitro*, the vectorized NT analog VH-N412 led to decreased NTSR2, as well as GFAP and Iba1 expression upon inflammatory conditions, suggesting that binding of VH-N412 on NTSR2 attenuates astrocytic and microglia reactivity during inflammation. It has been shown that activation of NTSR2 by its ligand can result in modulation of gene expression (Vita et al., 1998; Gendron et al., 2004; Ayala-Sarmiento et al., 2015). Inhibition of NTSR2 by JMV449, a general NTSR antagonist, suppressed GFAP expression, potentially by affecting transcriptional 3 (STAT3) factor signaling (Ando et al., 2019). Also, NT and NT agonist JMV449, significantly induced the proliferation rate of some astrocytic tumor cell lines, suggesting that NT is secreted by astrocytes and acts as an autocrine and/or paracrine growth factor (Camby et al., 1996). In our experimental settings we do not know whether VH-N412 activates or inhibits NTSR2 considering that different NTSR2 ligands such as NT, SR48692 (a selective non-peptide NTSR1 antagonist) and levocabastine, behave as agonists or antagonists, depending on specific signaling events and cell types studied (Vita et al., 1998; Dobner, 2005). Nevertheless, one may hypothesize that NTSR2 up-regulation during inflammation allows NT or its analogs to antagonize inflammatory processes and that NTSR2 could be a relevant pharmacological target in the modulation of neuroinflammation.

G protein coupled receptors (GPCRs) have been shown to play important roles in inflammation, and their involvement in transmembrane signaling is primarily responsible for the mediation of complex inflammatory (and anti-inflammatory) responses (reviewed in Sun and Ye, 2012). As such, the NT receptors and the neurotensinergic system have been involved in various inflammatory disease states. NT has been shown to promote an acute inflammatory response in an experimental model of colon inflammation (Castagliuolo et al., 1999). In contrast, an ameliorative effect of NT in the colitis model of inflammatory bowel disease has been reported, that reduces the levels of the proinflammatory cytokines IL-6 and tumor necrosis factor-α (TNF-α) that are highly expressed in this model (Akcan et al., 2008). Recently, NT exhibited anti-inflammatory activity in induced asthma, and this beneficial effect was mediated through NTSR1 (Russjan & Kaczyńska, 2019). With regard to brain diseases, NT has been studied in Schizophrenia, Alzheimer’s disease, and Parkinson’s disease among others (reviewed in St-Gelais et al., 2006). However, the specific role of NT in neuroinflammation in these disorders has not been addressed, nor which receptors are involved. A large and mild decrease of NTSR2 and NT mRNA respectively has been described in the temporal lobe of Alzheimer’s disease patients (Gahete et al., 2010).

### NTSR2 is expressed in blood vessel endothelial cells and increased in rodent models of epilepsy

Besides NTSR2 expression in astrocytes of the hippocampal parenchyma, we observed perivascular NTSR2 labeling that was significantly enhanced (an average of 3-fold increase) in the PILO rats in the hours and days that follow SE as exemplified at 3 days post-SE. Detailed studies addressing differences between parenchymal and perivascular astrocytes are lacking, and, although they are bound to have distinct functions, both showed increased GFAP and NTSR2 in the PILO rats. Confocal analysis of GFAP and NTSR2 double labeling immunohistochemistry showed some NTSR2 and GFAP co-localization in astrocytic end-feet in the PILO rats, that are polarized around the basement membrane to maintain the structural integrity of the BBB and restrict vascular permeability (Abbott et al., 2006). In addition, the area that surrounds the blood vessels in PILO rats contained a large number of reactive hypertrophic astrocytes known to be associated with neurologic disease, including inflammation and neurodegeneration (Hol & Pekny, 2015).

A number of molecules have been identified whose expression is increased in reactive astrocytes as well as in their perivascular processes. For instance, aquaporin-4 (AQP-4) that mediates the delivery of nutrients to surrounding neurons is increased in sclerotic hippocampi from patients with temporal lobe epilepsy and correlates positively with increased GFAP in the astrocyte plasma membranes, but also in perivascular end-feet (Lee et al., 2004).

One major finding of our study is that NTSR2 is not exclusively co-localized with GFAP but is also found distributed between astrocytic end-feet and blood vessel lumen. Triple immunolabeling experiments to localize NTSR2, together with the CD31, PDGFRβ and Collagen IV markers that specifically label endothelial cells, pericytes and basal lamina respectively, led to the finding that in addition to reactive perivascular astrocytes and their end-feet, NTSR2 is expressed in brain endothelial cells, and only faintly in pericytes in the PILO-treated rats. Co-upregulation of specific proteins in both astrocytes and endothelial cells in disease or inflammation has been reported. For instance, IL1β up-regulates GLUT1 both in astrocytes and endothelial cells (Jurcovicova, 2014). Similarly, Lcn2, a secreted lipophilic protein was strongly induced after LPS injury, not only in astrocyte reactive gliosis but also in endothelial cells (Zamanian et al., 2012). Taken together, our results indicate that NTSR2 is expressed by different components of the blood vessels, in particular perivascular astrocytes and endothelial cells in PILO rats and may be implicated in the structural integrity of the BBB. While human pulmonary artery endothelial cells (PAECs) were shown to express NTSR3 (Shults et al., 2018), to our knowledge, our study is the first to show NTSR2 expression in blood vessels in general and in the BBB in particular. This finding suggests that while NTSR2 induction is associated with neuroinflammation involving parenchymal and perivascular astrocyte reactivity, NTSR2 may also play a specific role in brain endothelial cells. For instance, NTSR2 could be involved in modulating endothelial cell properties due to its function as a signal transduction receptor. Indeed, different studies report that NT stimulation of NTSRs activates the extracellular signal regulated kinases 1/2 (ERK1/2) (Martin et al., 2003; Navarro et al., 2006; Sarret et al., 2002). Interestingly, it has been shown that NTSR2 is internalized after NT binding and that receptor internalization is prevented by the NTSR2 specific antagonist levocabastine (Nouel et al., 1999; Ayala-Sarmiento et al., 2015). Like for other GPCRs, receptor internalization plays critical roles in the signaling pathway (Gendron et al., 2004). Finally, it has also been shown that the mouse NTSR2 recycles to the cell membrane (Martin et al., 2002). While nothing is known at this stage for NTSR2 internalization and recycling in brain endothelial cells, this receptor could be involved in NT endocytosis, intracellular release of NT and recycling at the cell membrane, meeting the conditions for NT transcytosis across the BBB if these functional properties were confirmed in brain endothelial cells. The limited effects of NT on hypothermia, pain modulation, and behavior when administered i.v. compared to i.c. administration, are related at least in part to poor blood stability and low half-life. Stable NT analogues appear to elicit central effects when administered i.v. (Kokko et al., 2005; reviewed in Boules et al., 2006), suggesting trans-BBB transport. The potential of the VH-N412 vectorized NT to alleviate astrocyte reactivity *in vitro* suggests that peripheral NT transported across the BBB or NT endogenously produced in the CNS could modulate neuro-inflammation.

### NTSR2 is expressed as immediate early gene in glial cells stimulated with proinflammatory factors

The transient and rapid upregulation of NTSR2 and NTSR3 mRNA after treatment with proinflammatory agents such as LPS and IL1β led us to question whether these NTSRs could be categorized as IEGs. Conditions for IEG classification are that the gene’s mRNA levels are rapidly -in general within an hour- and transiently increased, in response to different cell stimuli. In addition, this upregulation should be independent of *de novo* protein synthesis (Mehendale & Apte, 2009; Fowler et al., 2011; Bahrami & Drabløs, 2016). Our results show that the NTSR2 and NTSR3 mRNA expression levels that were increased as early as 1 h after LPS were unaltered by CHX. The inflammation mechanisms or pathways involved in rapid induction of NTSR2 and NTSR3 expression remain unknown. It will be therefore essential to dissect and understand the properties of these new IEGs, how they are activated and modulated in response to biological stimuli, and how they impact downstream molecular processes and biological responses. It has been reported that several extracellular signals can activate different cellular pathways that upregulate transcription factors and IEGs. Depending on the cellular context and experimental conditions, the RhoA-actin, ERK and p38 MAPK and PI3K pathways have been implicated in the activation of transcription factors, leading to induction of IEGs (Fowler et al., 2011; Bahrami & Drabløs, 2016). One may hypothesize that NTSR2 and NTSR3 mRNA rapid induction in inflamed glial cells is mediated via one or several of these reported pathways.

IEGs encode different groups of genes including transcription factors, growth factors, cytoskeleton proteins, and receptors such as the NR4A subfamily of nuclear receptors (NR4A1, NR4A2 and NR4A3 (Martínez-González & Badimon, 2005; Mehendale & Apte, 2009; Zhao & Bruemmer, 2010). In line with our data, several studies revealed that the NR4A subfamily responds to inflammation (Maxwell & Muscat, 2006), similar to NTSR2 and NTSR3. The NR4A1, NR4A2 and NR4A3 are rapidly upregulated in macrophages after LPS stimulation (Barish et al., 2005; Pei et al., 2005; Maxwell & Muscat, 2006), and NR4A1 is also induced by several cytokines (Pei et al., 2005; Maxwell & Muscat, 2006). In addition, there is evidence for a relationship between the NR4A upregulation and the Nuclear factor NF-κB, one of the major mediators involved in the inflammatory pathway (Pei et al., 2005; Maxwell & Muscat, 2006). Further studies will be required to decipher the links between the immediate early responses of NTSR2 and NTSR3 and astroglial inflammation.

It is well established that IEGs play important roles in cell growth, differentiation, immune system responses, learning and memory long-term potentiation, inflammation, vascular reorganization and also in diseases (Davis et al., 2003; Plath et al., 2006; Mehendale & Apte, 2009; Bahrami & Drabløs, 2016). In the immune system several IEGs have been shown to play important roles in transmitting the activation signal downstream via transcription factors to dictate expression patterns of particular target genes with immunological significance. It is the case for example for Zif-268/egr-1 shown to play a role in immune responses by regulating downstream gene targets encoding interleukin-2 (IL-2), CD44, ICAM-1, tumor necrosis factor (TNF) and NR4A1 in immune cells such as macrophages, B and T lymphocytes (McMahon & Monroe, 1996; Bahrami & Drabløs, 2016). Microglia and astroglia can be considered as part of the CNS immune system that respond to and release various proinflammatory factors such as cytokines and chemokines (Lokensgard et al., 2002). Interestingly, in addition to the induction of NTSR2 and NTSR3 IEGs, our data showed that the transcription factor Zif-268/egr-1 IEG is also rapidly induced in glial cells treated with LPS, similar to observations in activated immune cells (Bahrami & Drabløs, 2016). Furthermore, in our study, several other inflammatory markers including GFAP, MCP1/ccl2, but also Iba1 and IL1β (data not shown) were also induced by LPS in activated glial cells. It is appealing to suggest that Zif-268/egr-1 may play a role in the triggering of the immune response in activated glial cells and that the induction of Zif-268/egr-1 involves pathways similar to those -or is involved in the pathway-leading to the expression of NTSR2, NTSR3 and other immune response factors mentioned above.

## CONCLUSION

In all, our work demonstrates the involvement of the neurotensinergic system in the parenchymal glia and glio-vascular unit in conditions of neurological diseases. Our results show that NTSR2 is expressed as an IEG in glial cells in response to proinflammatory factors and is implicated in astroglial and perivascular inflammation. An NT analog down regulates glial inflammation suggesting that NTSR2 may regulate glial responses to injury and that targeting the NTSR2 receptor may open new avenues in the regulation of neuroinflammation.

## ACKNOWLEDGEMENTS

This work was supported by a fellowship to GK from the European Union’s Horizon 2020 research and innovation program under the Marie Sklodowska-Curie grant agreement No 642881, and AMIDEX (ICN PhD Program, ANR-11-IDEX-0001-02 grant) funded by the French Government «Investissements d’Avenir» program. Funding from the CNRS and Aix-Marseille Université (AMU) to the Institute of Neurophysiopathology (INP), UMR7051 is acknowledged. We thank Fanny Gassiot for her expertise in VH-N412 synthesis and Antoine Ghestem for advice on the rat pilocarpine model. We are grateful to Dr. Jean-Pierre Kessler and Dr. Fabien Tell for helpful discussions and sharing lab space and equipment. We also thank Stéphane Girard, Christophe Fraisier, and Yasmine Mechioukhi (all from Vect-Horus SAS, Marseille) for providing PDGFRβ, CD31 and ColIV antibodies. Last, we thank Dr. Mourad Mekaouche and Axel Fernandez for advice and help with animal care, as well Yves Gobin for laboratory assistance.

## FIGURE LEGENDS

**Figure S1.**
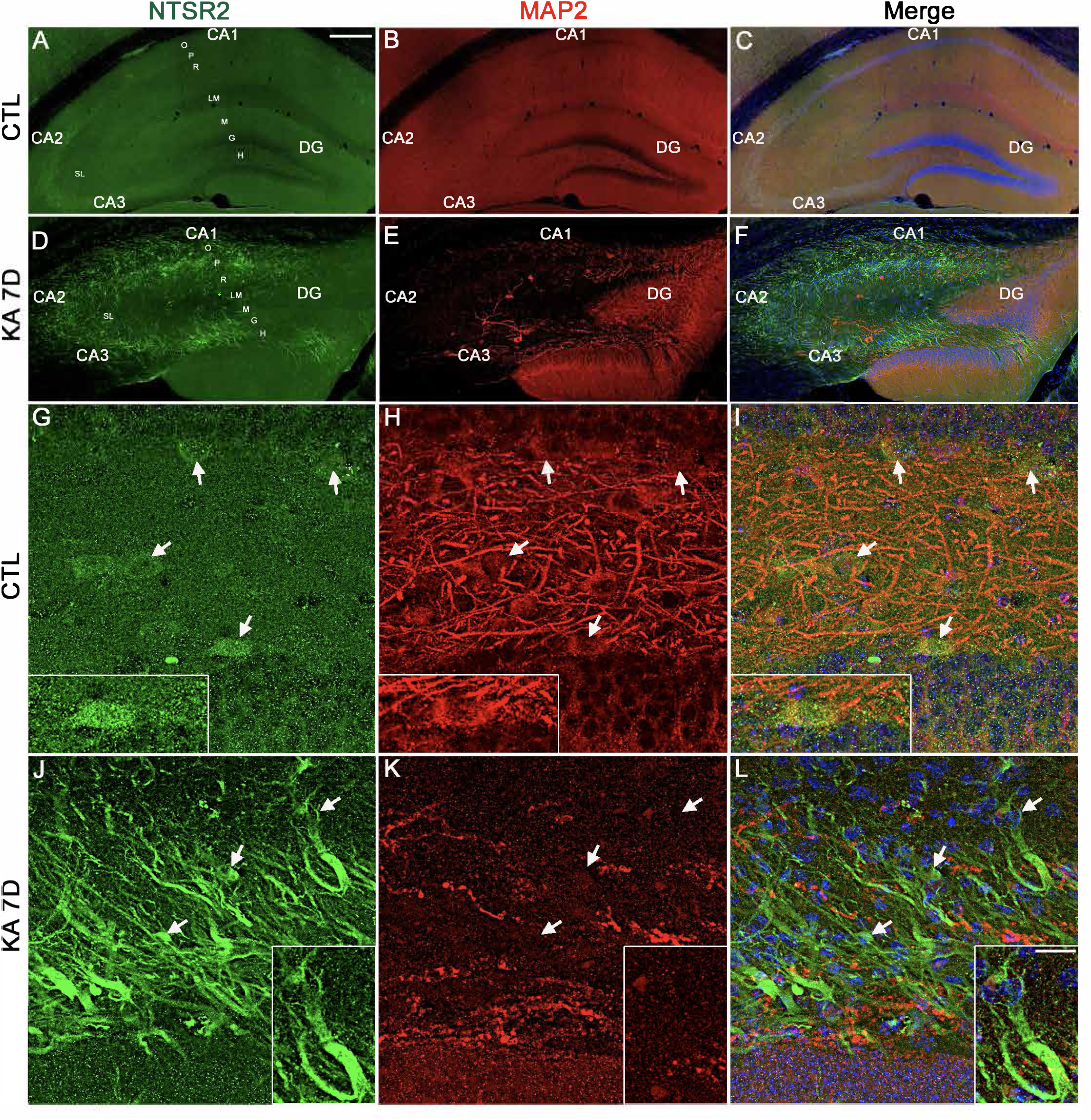
NTSR2 expression is increased in mice hippocampus following KA-induced SE. NTSR2 (green) and MAP2 (red) double immunolabeling was performed in control (CTL, A-C, G-I) and KA-treated mice at day 7 post-SE (KA 7D, D-F, J-L). High magnifications of the hilus of CTL (G-I) and KA 7D (J-L) are shown. In CTL, NTSR2 immunolabeling is clearly shown in neurons, identified by MAP2 staining (see arrows and insets). In KA 7D, NTSR2 immunolabeling is strongly increased in cells that are not immunolabeled for MAP2, compared to CTL mice. NTSR2 immunoreactivity was observed in non-neuronal cell bodies and processes (see insets), likely astrocytes. Panels C, F, I and L correspond to superimposition of NTSR2, MAP2 and DAPI. Cell nuclei were counterstained with DAPI (blue, C,F,I,L) that underlined the different areas of the hippocampus and its deformation and degeneration that occur in the KA model of epilepsy. Scale bars: 450 μm in A-F, 10 μm in G-L and 5 μm in insets. O: stratum oriens; P: pyramidal neurons of CA1, CA2 and CA3; R: stratum radiatum; LM: stratum lacunosum-moleculare; M: molecular layer; DG: dentate gyrus; G granule cells of DG; H: hilus of DG.

**Figure S2.**
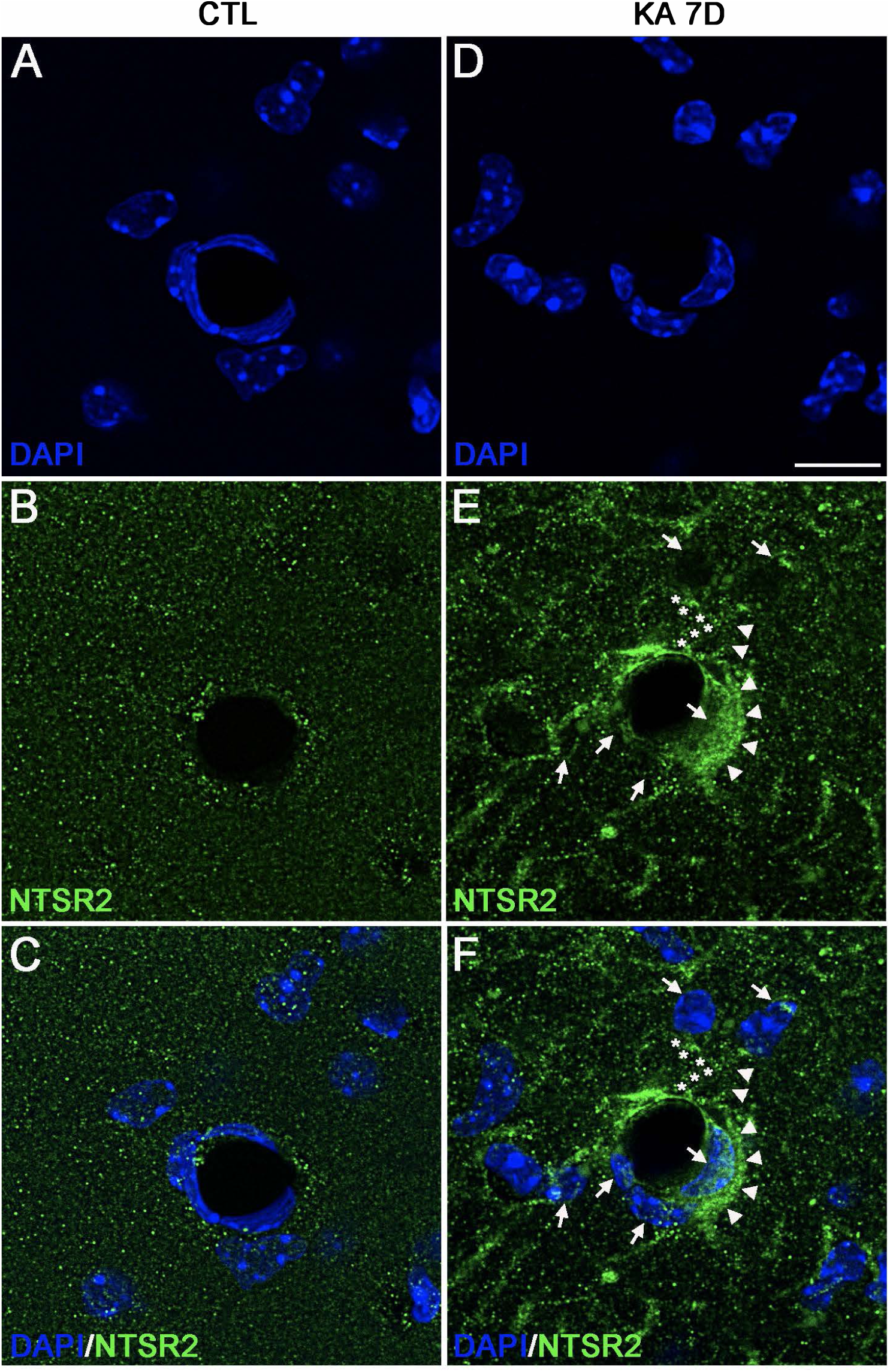
NTSR2 expression is increased in mice blood vessels following KA-induced SE. NTSR2 (green) immunostaining was performed in CTL (A-C) and KA-treated mice at day 7 post-SE (KA 7D, D-F). In KA 7D, NTSR2 immunoreactivity was increased in blood vessel cells of the lacunosum moleculare (LM) compared to CTL mice. In addition, several cells around these vessels displayed moderate to strong NTSR2 expression compared to CTL mice (arrowheads). Other dispersed cells in LM expressing NTSR2 were distinguished, projecting their processes (indicated by arrows and asterisks) around the blood vessels, similar to observations in blood vessels of pilocarpine rats. Cell nuclei were counterstained with DAPI (blue, A,C,D,F). Scale bar: 20 μm in all panels.

## Notes

**CONFLICT OF INTERESTS STATEMENT** MK is director of the Institute of Neurophysiopathology, UMR7051, an academic neuroscience laboratory supported by the CNRS and Aix-Marseille University, but also co-founder, shareholder and scientific counsel of the VECT-HORUS biotechnology company.

### Competing Interest Statement

MK is director of the Institute of Neurophysiopathology, UMR7051, an academic neuroscience laboratory supported by the CNRS and Aix-Marseille University, but also co-founder, shareholder and scientific counsel of the VECT-HORUS biotechnology company.

